# A Realistic Simulation Framework for Evaluating Microbiome Normalization in Sample Stratification and Differential Abundance

**DOI:** 10.64898/2026.01.13.699216

**Authors:** Amen Al Khafaji, Carolina Gómez-Llorente, José Camacho

## Abstract

**Background:** Normalization is a critical yet often poorly understood step in microbiome studies. Suboptimal approaches may lead to inaccurate conclusions in downstream analyses of microbial communities. Currently, there is no benchmarking framework to evaluate how normalisation affects both sample stratification and differential abundance simultaneously across taxonomic levels. In this paper, we propose a simulation pipeline based on real data and multivariate exploratory data analysis to provide a structured and reproducible assessment of normalization methods.

**Results:** Normalization methods exhibited distinct accuracy across taxonomic levels and sequencing depths. In our case study, at the phylum level, edgeR-TMM and Rarefaction improved accuracy by reducing coverage-related variation while preserving biological structure. In contrast, at the genus level, the overall improvement by normalization was less pronounced, reflecting the weaker influence of sequencing depth variability in this scenario, and EdgeR-TMM again provided the most accurate estimation of biological effect. Multivariate visualizations supported these observations, highlighting both sample-level and taxon-level differences among methods. Yet, ordination-based summaries are not sufficient for differential abundance inference and can be misleading, motivating the use of a simulation environment with known ground truth.

**Conclusions:** Normalization performance varied with sequencing depth, sparsity, taxonomic resolution, and dataset size. Thus, there is no single normalization method that is expected to be optimal across all conditions. Our proposed simulation and analysis framework offers a reproducible and interpretable platform to evaluate existing and new normalization approaches in microbiome research for specific case studies.

## 1 Background

Understanding human microbiome composition helps us to understand how specific microbes interact with disease, where an alteration of this composition occurs, which is called dysbiosis. The study of the human microbiome composition allows the development of personalized diagnostic and therapy decisions [1–3]. Metagenomic studies capture these interactions by analysing the abundance and diversity of microbial species in a sample using high-throughput DNA sequencing [3]. To profile such communities, researchers use two sequencing approaches: shotgun sequencing and marker gene analysis [4–6]. In shotgun sequencing, all microbial genes in a sample are sequenced, whereas marker gene analysis focuses on a single phylogenetically informative gene, typically 16s or 18s rRNA [7, 8]. The 16s rRNA gene is the widely accepted method when the interest lies in the specific microbial composition of a sample [7, 9, 10]. Following sequencing, reads are processed to generate a taxa abundance table, representing counts of each taxonomic group across samples. These taxonomic units are defined as either Operational Taxonomic Units (OTUs), obtained through clustering, or Amplicon Sequence Variants (ASVs), derived from error-correction methods [11–13]. The sum of counts for each sample, known as its sequencing depth or library size, varies considerably between samples [12]. Metagenomic datasets, like other omics data, are affected by both biological and technical sources of variation. Among these, differences in sequencing depth across samples represent a major challenge, affecting the accuracy of downstream analyses [12]. Normalisation is therefore essential to ensure comparability between samples. Yet, there is no consensus on the best way to normalize these data.

A common normalization approach is Total Sum Scaling (TSS) [14], in which all samples are scaled to the same sum of counts [15, 16]. Rarefaction, also called subsampling, randomly resamples reads from each sample to a fixed sequencing depth, which may lead to the loss of rare or low-abundance species and reduce statistical power [11, 12, 16–20]. Beyond these approaches,count-based statistical methods originally developed for transcriptomics, such as DESeq [21–23] and edgeR [21, 22, 24], provide alternatives that account for differences in sequencing depth without discarding data. Library-size normalization adjusts for differences in sequencing depth but does not account for the compositional nature of microbiome data, where counts are constrained by a fixed total. Analyses must therefore focus on relative abundances rather than absolute counts [17]. Compositional Data Analysis (CoDA) provides statistical methods designed for such data [25–27]. A common approach is the Centered Log-Ratio (CLR) transformation, in which each taxon’s count is divided by the geometric mean of all taxa in a sample and log-transformed [27, 28]. The choice of an optimal normalization method remains challenging, and recommendations in different studies are often in conflict [5, 18, 25, 29, 30], as shown in Table 1. Existing evaluations typically focus on a single analytical aspect, either sample-level (stratification or clustering) or feature-level (differential abundance). Rarely do they assess both within an integrated framework. This narrow focus provides only a partial view of the performance of the method. Another important issue is that studies using real data lack ground truth (the actual difference in abundance between two ecosystems), making it difficult to evaluate [31, 32]. Similarly, many simulations may not fully capture the compositional nature of microbiome data, limiting interpretability and reliability.

**Table 1:** Assessment approaches applied in previous studies.

| Study (Data type) | Focus of the study with and recommendation on normalization |
| --- | --- |
| McMurdie & Holmes (2014) [18] (Simulation) | Focus: Clustering and differential abundance (two separated simulations).<br>Recommendation: Against rarefying and proportions; mixture models perform better (visualization: PCoA).<br>Impact in the Field in Total References: 2871; Annual Rate: 280. |
| Weiss et al. (2017) [25] (Real data & Simulation) | Focus: Clustering and differential abundance (revisits [18] with real data).<br>Recommendation: Rarefying helps with depth variability. Against proportions (visualization: PCoA).<br>Impact in the Field in Total References: 1700; Annual Rate: 250. |
| McKnight et al. (2019) [34] (Real data & Simulation) | Focus: Clustering accuracy.<br>Recommendation: For proportions and rarefaction; against log transformations (visualization: PCoA).<br>Impact in the Field in Total References: 301; Annual Rate: 70. |
| Nearing et al. (2022) [29] (Real data) | Focus: Differential abundance.<br>Recommendation: For compositional tools (ALDEx2, ANCOM-II); against edgeR. Consensus analysis across methods advised (visualization: PCoA).<br>Impact in the Field in Total References: 340; Annual Rate: 170. |
| Schloss (2024) [30] (Simulation) | Focus: Clustering accuracy.<br>Recommendation: Rarefaction is the best current method for uneven sequencing effort (visualization: GGplot2).<br>Impact in the Field in Total References: 24; Annual Rate: 24. |

This summary highlights how the focus of analysis (e.g., clustering vs. differential abundance) influences recommendations on normalization methods. Both total references and annual rates were collected from Google Scholar according to the date of manuscript submission. Abbreviations: ANCOM = Analysis of Composition of Microbiomes; PCoA = Principal Coordinates Analysis.

To address these gaps, we introduce a realistic simulation framework, starting from real measurements, which incorporates a multivariate analytical approach based on Principal Component Analysis - observation-based Missing data methods for Exploratory Data Analysis (PCA–oMEDA) [33]. By embedding PCA–oMEDA within a ground-truth simulation pipeline, the framework provides a unified platform to evaluate normalization methods. Therefore, our study contributes in two major ways:

- Simulation-based evaluation platform: We embed PCA–oMEDA within a Monte Carlo simulation pipeline that generates datasets starting from real microbiome data to establish a known ground truth. The framework is not intended to identify a single “best” method, a goal complicated by the context-dependent nature of microbiome data. Instead, it provides a general framework for evaluating existing and new normalization methods across different scenarios within a single, coherent workflow.
- Integrated analytical approach: Previous studies relied on Principal Coordinates Analysis (PCoA) to visualize sample clustering [5, 18, 25, 29, 30]. PCA–oMEDA not only take advantage of sample clusters, but also can simultaneously evaluate the accuracy in the identification of the taxa responsible for the differences.

## 2 Methods

### 2.1 Simulation and Analysis Workflow

The workflow in this study evaluates normalization methods in metagenomic data analysis through a structured process. It involves simulating abundance data, applying normalization, and analyzing the results with PCA and oMEDA. Finally, the outputs are compared with the ground truth (Fig. 1). To illustarte the workflow, we simulate abundance data derived from real microbiome data reported in the study “A MultiOmics Approach Reveals New Signatures in Obese Allergic Asthmatic Children” [35]. An abundance table **X** was generated, with rows representing individual samples and columns representing taxa (bacterial groups), and each sample labeled by condition (*healthy vs. dysbiosis*). Two simulations, one at phylum and other at genus level, were designed to capture the biological variation of bacteria in the human gut microbiome.

**Fig. 1:**
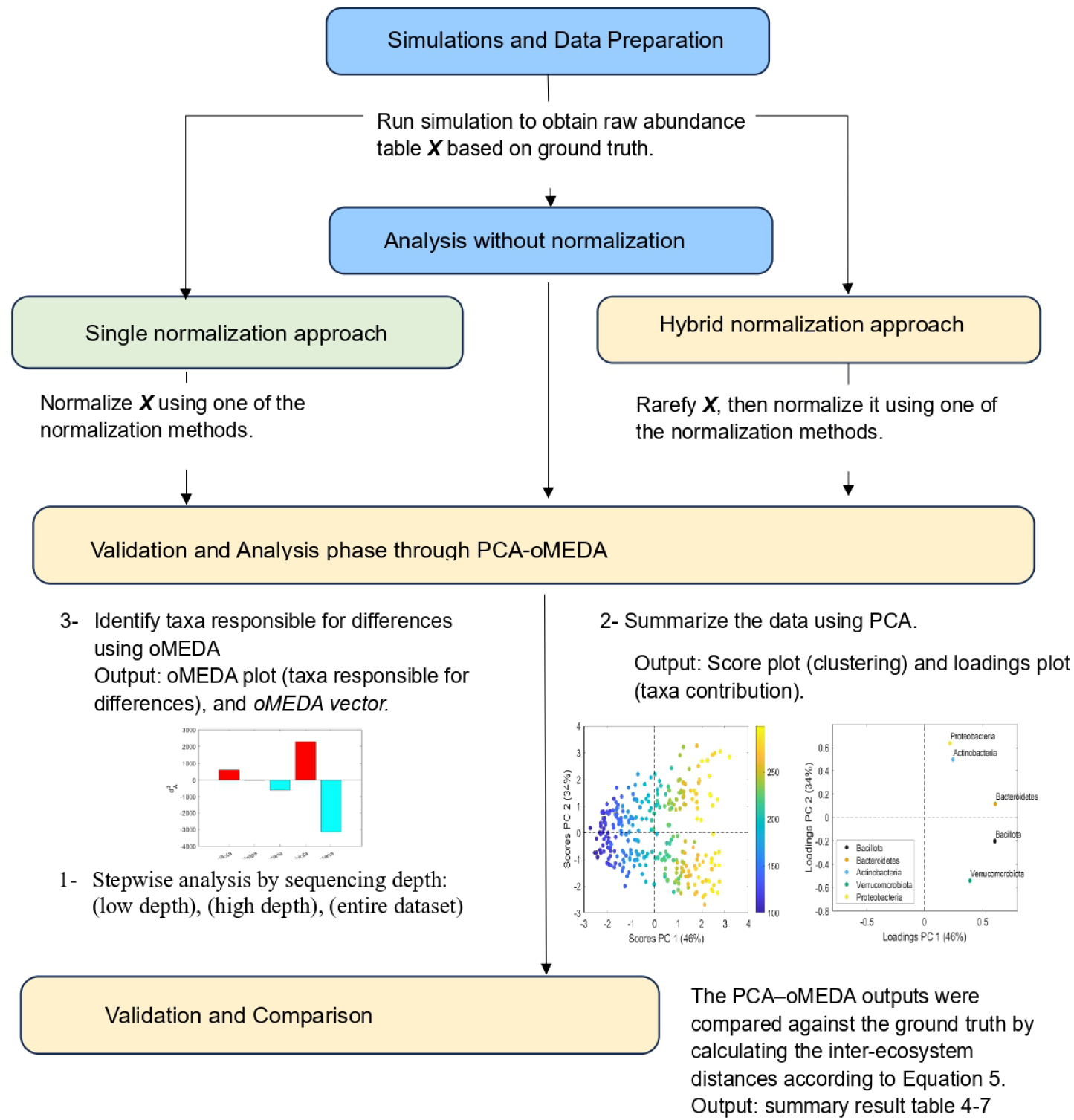
Workflow of the proposed method for evaluation of the normalization of metagenomic data. Abbreviations: PCA = Principal Component Analysis; oMEDA = observation-based Missing data methods for Exploratory Data Analysis.

These simulations were conducted at different taxonomic level to provide multiple scenarios, accounting for both the compositional nature and the sparsity of microbiome data. Because the composition of the simulated data is known, these datasets provide a controlled framework for evaluating the performance of different normalization methods. By systematically varying the parameters of these simulations, we were able to explore how different sequencing depths and taxonomic resolutions affect the ability of PCA and oMEDA to capture underlying biological differences. We first modeled a simulation at the phylum level, followed by a simulation at the genus level.

#### 2.1.1 Phylum-level simulation

The phylum-level simulation is created based on bacterial taxa at the phylum level, which are characterised by a minimal occurrence of zero reads. It includes five bacterial phyla: *Bacillota*, *Bacteriodota*, *Actinomycetota*, *Verrucomicrobiota*, and *Pseudomonadota* [35]. For each ecosystem, a pool of 10^6^ sequences was generated, with each phylum represented according to its target percentage, as illustrated in Fig. 2. From this pool, individual samples were created by randomly selecting between 100 and 300 reads with replacement. This sampling procedure was repeated to produce 150 samples per condition (healthy and dysbiosis). The resulting dataset provides a controlled framework to evaluate normalization methods and assess their performance in handling variable phylum abundances and low-frequency taxa.

**Fig. 2:**
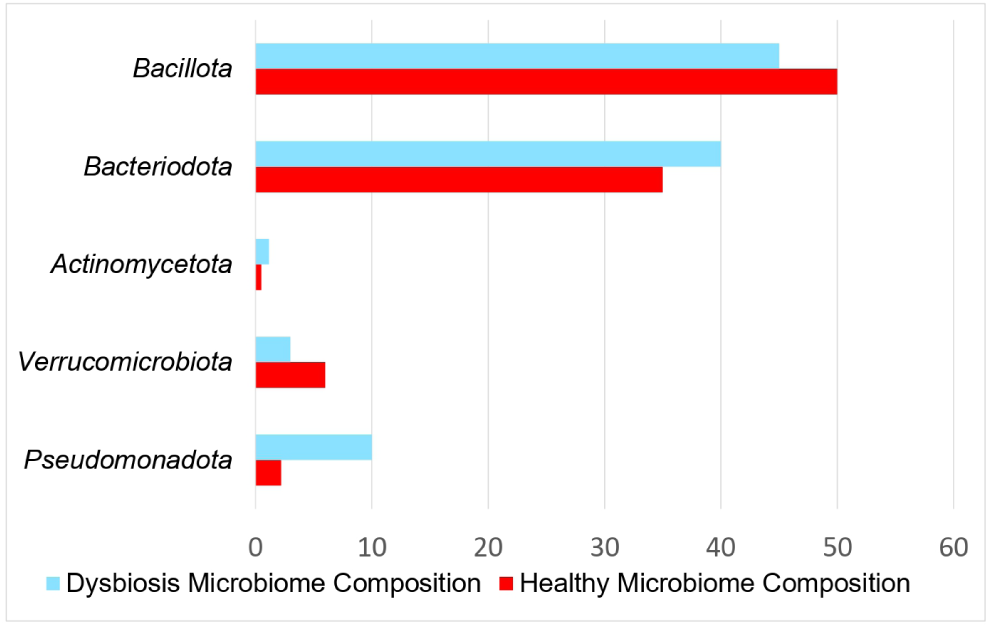
Illustration of the percentage (ground-truth) of each bacterium at phylum level.

#### 2.1.2 Genus-level simulation

The genus-level simulation was designed to assess normalization methods under conditions with a high proportion of zero counts, reflecting the higher diversity at the genus level. The simulation included 36 features (taxa), generated according to the percentages shown in Fig. 3. The simulation used the same sequence generation and sampling depth parameters as the phylum-level simulation. Each sample contained between 100 and 300 reads drawn randomly with replacement. Approximately 60% of samples had zero counts for rare taxa. Some normalization methods, such as CLR, cannot handle zeros. Zeros were thus replaced with a pseudo-count of 1, enabling log-transformations while preserving the overall compositional structure of the data.

**Fig. 3:**
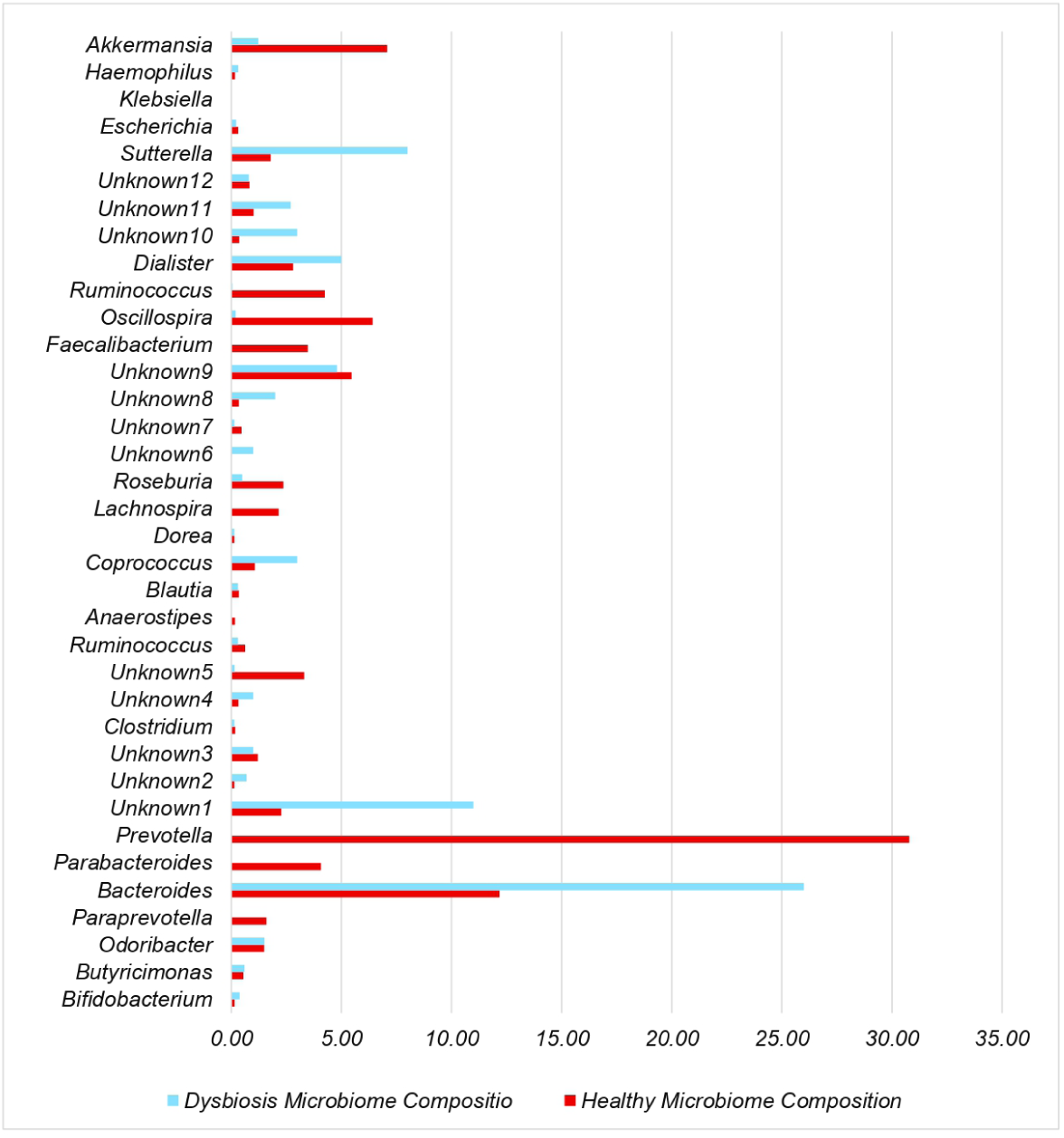
Illustration of the percentage (ground-truth) of each bacterium at the genus level.

### 2.2 Normalization

After creating abundance tables from the simulations, normalization methods were applied to correct for differences in sequencing depth across samples. Two approaches were considered: a direct normalization approach, where a method from Table 2 was applied directly to the raw abundance data, and a hybrid normalization approach proposed in this study.

**Table 2:** Normalization methods evaluated in this study.

| Name | Description |
| --- | --- |
| None | No correction for unequal sequencing depth. |
| Rarefaction | Subsample each sample to equal depth without replacement [25]. |
| Center Log Ratio | Log-transform values relative to the geometric mean of each sample [28]. |
| Total Sum Scaling | Divide taxa values by the sum of counts in each sample [28]. |
| Cumulative Sum Scaling | Scale counts using a stable quantile of each sample’s distribution, reducing the impact of high counts [28]. |
| DESeq2 | Computes scaling factors using geometric means and median ratios, stabilizing variance with a log-like transformation [25]. |
| edgeR-TMM | Computes trimmed mean of log-fold differences relative to a reference to derive scaling factors [24, 25, 28]. |

#### 2.2.1 Single normalization approach

In the Single normalization approach, a normalization method from Table 2 was applied directly to the raw abundance table. This includes methods that convert counts to relative abundances (e.g., TSS), estimate sample-specific scaling factors (e.g., CSS, DESeq2, edgeR-TMM), or transform the data to a log-ratio scale (e.g., CLR).

#### 2.2.2 Hybrid normalization approach

The hybrid approach proposed in this study combines rarefaction with a subsequent normalization step. Rarefaction is used to equalize sequencing depth across samples [18]. However, it does not in itself provide the additional scaling or transformation required by many downstream differential abundance workflows. Therefore, after rarefaction, a normalization method from Table 2 was applied to the resulting rarefied table in the same manner as in the direct approach. This hybrid approach standardizes sequencing depth prior to normalization while retaining the flexibility of applying different post-normalization procedures. However, it also inherits limitations of rarefying, including information loss due to subsampling and additional variability introduced by the random sampling step [18].

### 2.3 PCA and oMEDA Analysis

Multivariate methods allow the analysis and visualization of the abundance table **X** or data structures derived from it. In this study, we focus on PCA and oMEDA, which form the core of our pipeline. Throughout this paper, samples are arranged in the rows of **X**, while taxa are arranged in the columns, yielding *N* rows and *M* columns.

#### 2.3.1 Principal Component Analysis (PCA)

PCA is a well-known multivariate analysis method used to reduce the dimensionality of high-dimensional data while preserving most of its variance [27, 36, 37]. PCA generates a new representation of the data composed of orthogonal variables known as Principal Components (PCs), which capture the subspace containing maximum variance. In our pipeline, PCA is directly applied to the abundance table **X** after mean-centering. The PCA decomposition of matrix **X** is expressed as:

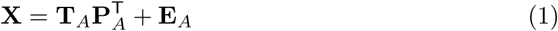

where **T***_A_*is the scores matrix (*N* × *A*), **P***_A_* is the loadings matrix (*M* × *A*), and **E***_A_* is the residual matrix (*N* × *M*).

PCA is similar to PCoA, also referred to as Classical Multidimensional Scaling (CMDS). While PCA operates directly on the data matrix, PCoA computes principal coordinates from a distance matrix (e.g., Euclidean, Bray–Curtis, or Jaccard) using eigen-decomposition. Notably, PCA on the mean-centered abundance table **X** is mathematically equivalent to PCoA on the corresponding Euclidean distance matrix. In this case, the PCA scores **T***_A_* coincide with the principal coordinates in PCoA for the same subspace dimension (*K* = *A*) [38, 39]. PCoA can accommodate non-Euclidean distances, allowing it to capture nonlinear relationships. However, this flexibility comes at the cost of reduced interpretability. A key advantage of PCA over PCoA is that the loadings **P***_A_* provide interpretable information about the taxa contributing to differences among samples. By using PCA, we retain the ability to visualize relationships among samples while simultaneously identifying the taxa driving these differences.

Fig. 4 compares samples from healthy and dysbiotic ecosystems using PCA on phylum-level simulations with no normalization. In Fig. 4a, the score plot shows the first two PCs. A color gradient indicates sequencing depth, which correlates with PC1 (horizontal axis, accounting for 46% of the variance). Fig. 4b displays the same plot colored by ecosystem, with healthy samples as black circles and dysbiotic samples as yellow circles. Separation occurs primarily along PC2 (vertical axis, accounting for 34% of the variance) and increases with sequencing depth. Thus, PCA shows that the confounding technical variability due to differences in sequencing depth is larger than the actual biological difference of interest (46% vs 34%). When combined with the score plot, the loading plot (Fig. 4c) provides information about how individual taxa connects to the main patterns of variance. We may interpret that *Bacteroidota* and *Pseudomonadota* are more abundant in dysbiotic samples, since loadings yield positive values in PC2, while *Bacillota* and *Verrucomicrobiota*, with negative loadings in PC2, are more abundant in healthy samples. Yet, the simulataneous interpretation of scores and loadings require some experience and can become challenging in some situations.

**Fig. 4:**
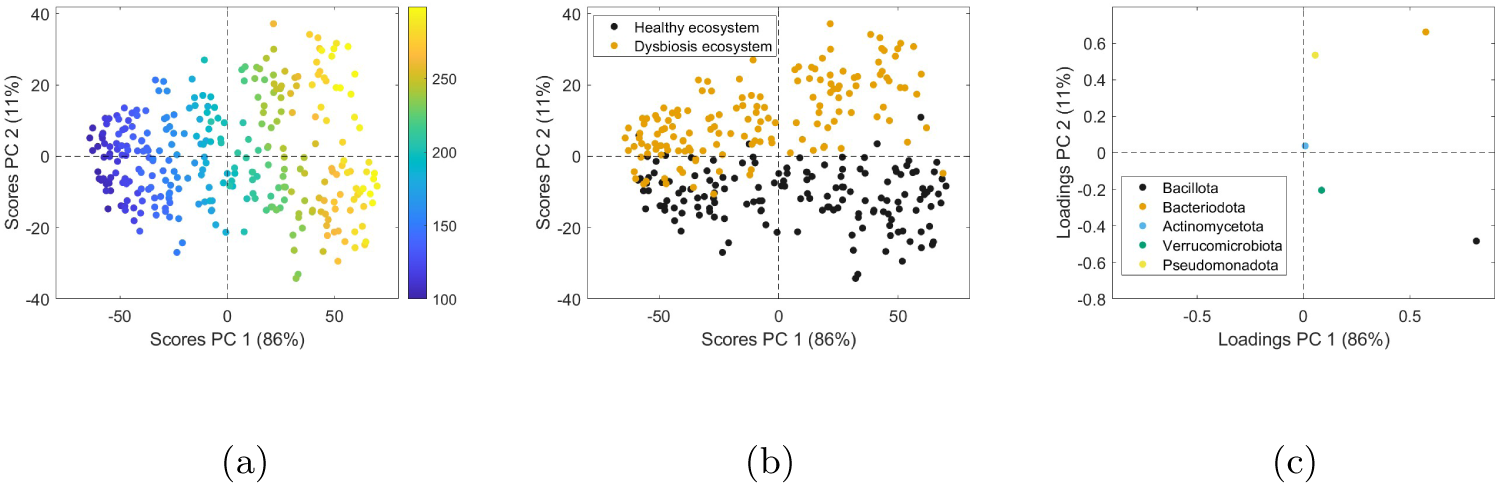
Score and loading plots of PCA for the data before normalization at the phylum level: **(a)** score plot of samples colored by sequencing depth, **(b)** score plot colored by the two ecosystems, and **(c)** loading plot of bacteria at the phylum level with five taxa.

The PCA procedure applied at the genus-level simulation is shown in Fig. 5. This maintains consistency in preprocessing and visualization with the previous analysis, allowing direct comparison at different taxonomic levels. The resulting score plot indicates increased separation between healthy and dysbiotic samples along PC1, explaining 80% of the variance, while differences in sequencing depth are mainly found in PC2, with 10% of the variance. This is a more favorable situation than the one depicted for phylum-level data, but is also challenging due to the number of taxa involved and the increased probability of 0 values. The corresponding loading plot identifies species-level contributions, with *Prevotella* (at the far right of PC1) enriched in healthy samples and *Sutterella* (at the far left of PC 1) enriched in dysbiosis, highlighting the taxa driving ecosystem differences.

**Fig. 5:**
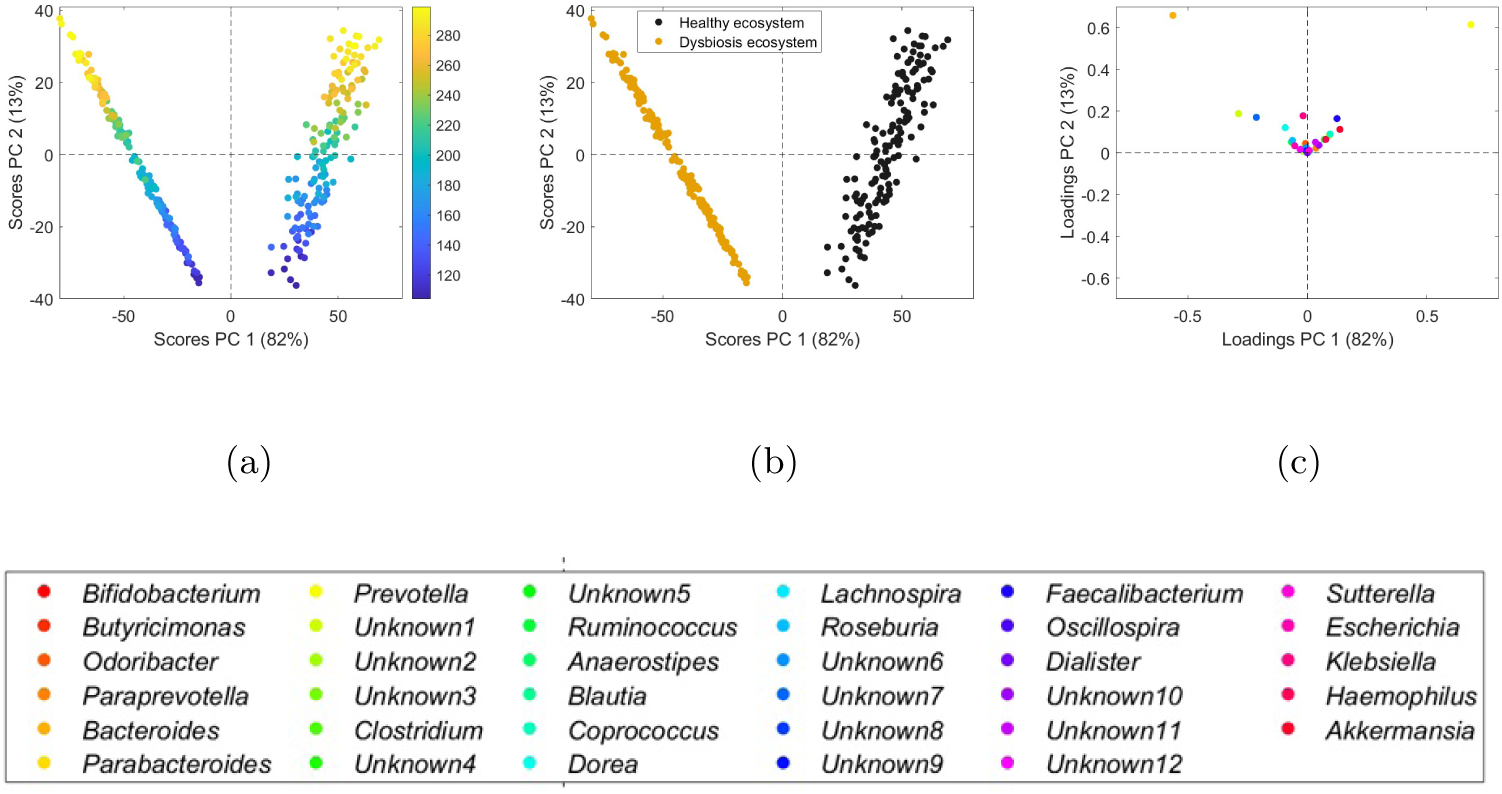
Score and loading plots of PCA for the data before normalization at the genus level: **(a)** score plot of samples colored by sequencing depth, **(b)** score plot colored by the two ecosystems, and **(c)** loading plot of bacteria at the genus level with 36 taxa. The full-width panel below shows the overall summary figure.

#### 2.3.2 Observation-based Missing data methods for Exploratory Data Analysis (oMEDA)

oMEDA [33] is an exploratory tool designed to facilitate the interpretation of relationships between the scores (**T***_A_*) and loadings (**P***_A_*) in multivariate models, such as PCA and Partial Least Squares (PLS) [37, 40]. It identifies variables contributing to specific patterns in the observations, including clusters, trends, or outliers. For that, the input to oMEDA is the multivariate model along with a specific numerical coding of the observations, with the coding describing the pattern of interest. In the case of this paper, we use a coding associated to the two ecosystems, with healthy samples identified by code 1 and dysbiosis by code -1. This coding allows us to understand the relative relevance of the loadings (i.e., the taxa) to the averaged separation between the scores of the two ecosystems in the PCA model. The output of oMEDA is a vector with that relative relevance, where positive values identify enriched taxa for healthy samples and negative values for dysbiosis samples. This can also be visualized as a bar plot that we call the oMEDA plot. The method is available through the MEDA toolbox [41].

To illustrate how oMEDA simplifies the interpretation of PCA, we applied it to phylum-level and genus-level data without normalization, in order to identify taxa driving differences between healthy and dysbiotic ecosystems. At the phylum level, Fig. 6, the oMEDA plot highlights taxa associated with each ecosystem. The analysis on non-normalized data only allows identifying the association of *Bacillota* (with a large negative bar) to healthy samples (code -1), and of *Pseudomonadota* (with a large positive bar) to dysbiosis (code 1). If we compare this result to the ground-truth in Fig. 6(b), we can see that oMEDA is only partially right. The mismatch between oMEDA and the ground-truth is the result of the confounding influence of the technical variability in the non-normalized data. Baring this idea in mind, we can assess the quality of a normalization method by analyzing the resulting data with PCA, and comparing the oMEDA result with the ground-truth. This comparison combines the accuracy in the clustering of samples (in our case, how well healthy samples are differentiated from dysbiosis samples in PCA) with the accuracy of the differential abundance (how well taxa associated to the biological effect are identified).

**Fig. 6:**
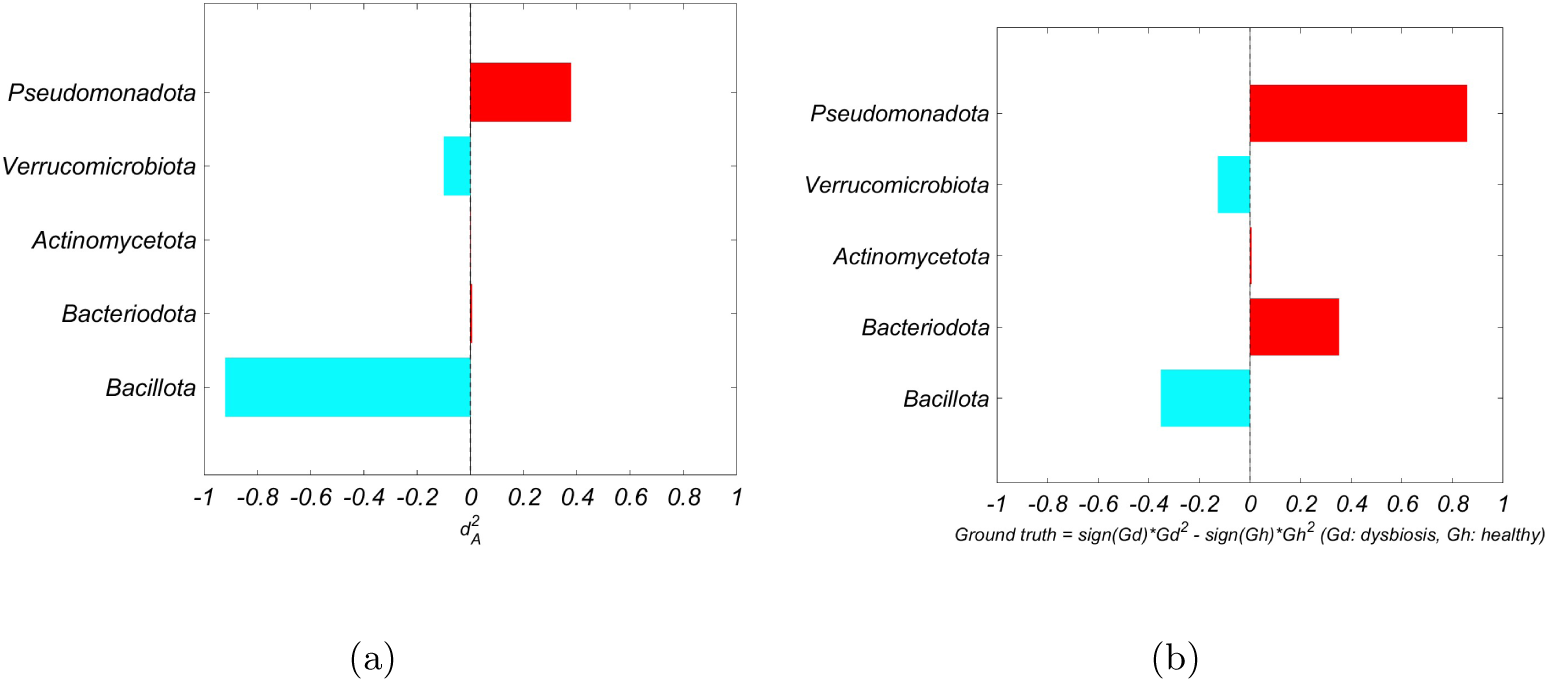
Score and comparison plots at the phylum level: **(a)** Observation-based missing data methods for exploratory data analysis (oMEDA) representation of raw data without normalization. **(b)** Ground-truth computed as the difference in bacterial composition between healthy and dysbiosis at the phylum level.

The same comparison is repeated at genus level (Fig. 7a vs 7b). In this case, main taxa correlated to the biological effect are mostly identified. Recall from Fig. 5 that this second simulation is more favorable in terms of the amount of biological variance in comparison to the technical variance, and this is reflected in the oMEDA accuracy. Yet, there is still a lot of information missing about low-abundance differences. Normalization may improve this situation.

**Fig. 7:**
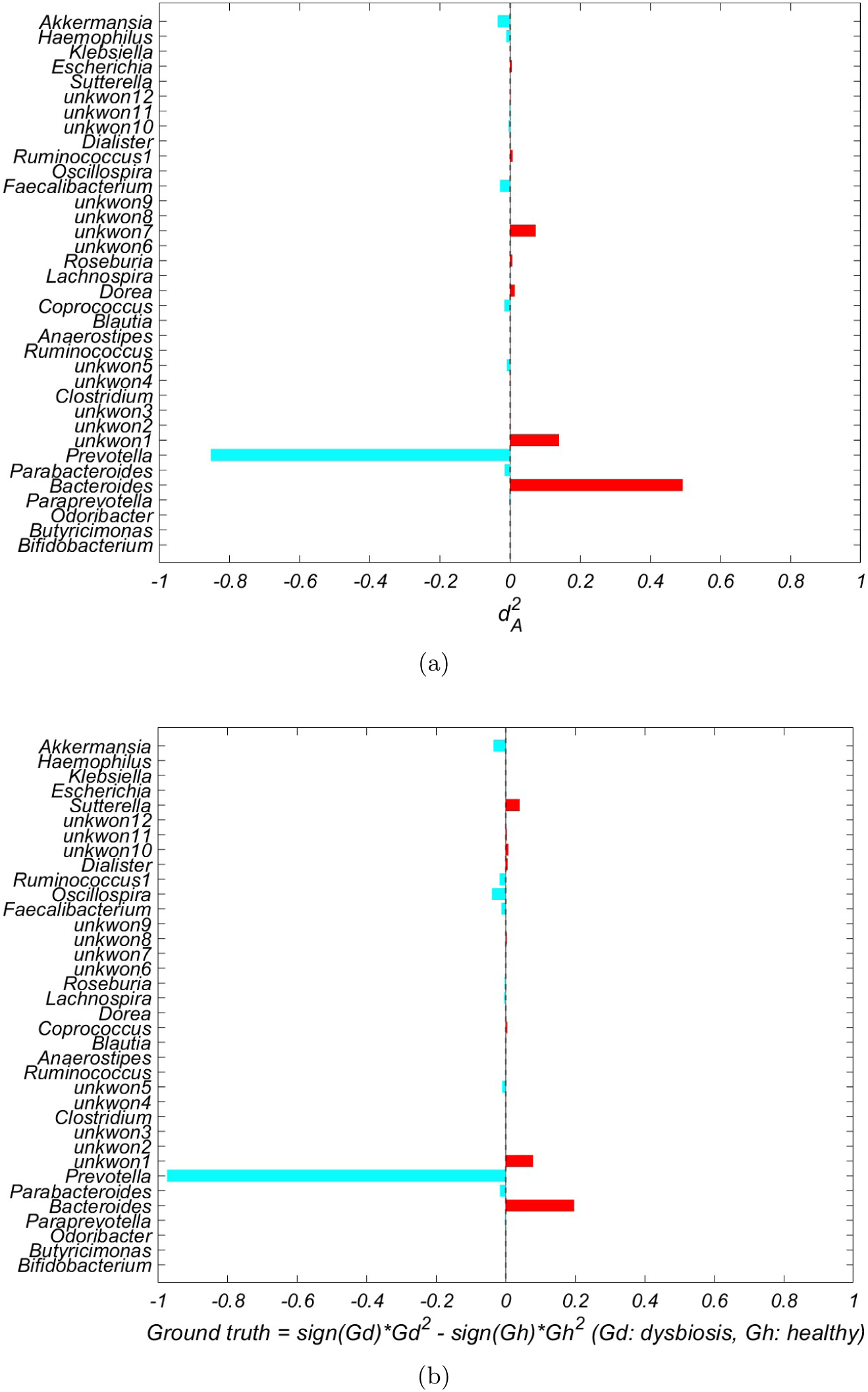
(**a**) Observation-based missing data methods for exploratory data analysis (oMEDA) representation at the genus level of raw data without normalization. (**b**) Ground truth computed as the difference in bacterial composition between healthy and dysbiosis at the genus level [35].

The role of oMEDA in this paper is to facilitate the interpretation of the connection between an effect of interest in the score plot (the difference between healthy and dysbiosis ecosystems) and the associated taxa connected to such effect. The oMEDA vector is also a numerical structure that can be easily compared to the ground-truth, and error measures between both of them can be easily computed.

### 2.4 Performance Benchmarking Against the Reference Standard

For the phylum simulation, Table 3 illustrates the ground truth. Positive values indicate taxa more abundant in dysbiosis samples, whereas negative values indicate taxa enriched in healthy samples.

**Table 3:**
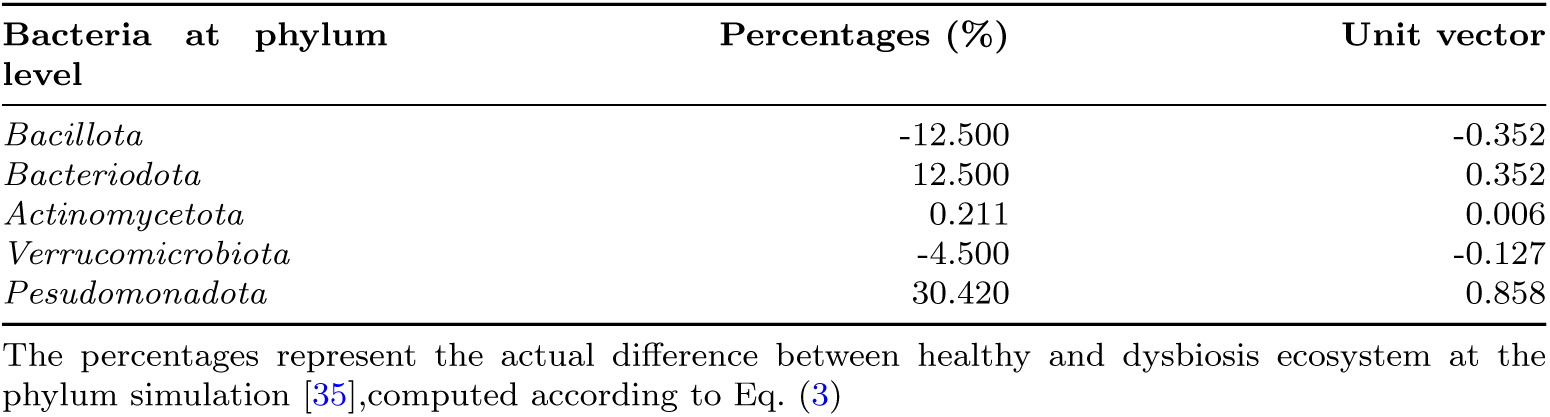
The groundtruth of phylum simulation.

Since oMEDA is based on squared distances between the two conditions, we computed the ground truth using squared, sign-preserving differences. This step allows a direct comparison between the oMEDA results and the ground truth. Mean-centering is also applied in the oMEDA setting; accordingly, taxa abundances were first mean-centered across the healthy and dysbiosis conditions when constructing the ground truth Eq. (2) before computing the between-group contrast Eq. (3).

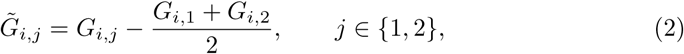

where *G_i,_*_1_ is the percentage of taxon *i* in healthy samples and *G_i,_*_2_ is the percentage of taxon *i* in dysbiosis samples.

Then, the squared (sign-preserving) ground-truth contrast was computed as:

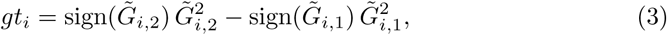

where *gt_i_* denotes the ground-truth difference for taxon *i*. Subsequently, both the ground truth vector and the corresponding oMEDA output vector were scaled to unit length to ensure comparability.

The performance of each normalization method was quantified using an error metric defined as the squared deviation between the oMEDA output and the ground truth (both scaled to unit length). Because oMEDA may flip sign arbitrarily, the error was evaluated for both possible directions of the ground truth and the minimum value was retained:

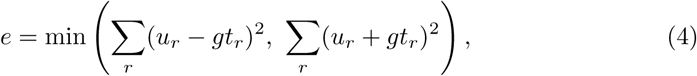

where *u_r_* represents the oMEDA differences between groups (for any normalization approach described in Section 3), and *gt_r_* denotes the ground truth differences. The normalization method achieving the lowest error was considered the best fit for the simulated dataset. Same settings applied at genus simulation.

## 3 Results

Table 4 shows the estimated error ratios across normalization methods for the phylum simulation. Results are reported separately for the subset of samples with 100–200 reads (referred to as low-depth samples), 200–300 reads (referred to as high-depth samples), and the complete dataset. The raw data results in the first row of the table indicate that the error at low depth is almost twice the error at high depth. When the complete dataset is considered, the maximum error is obtained. This pattern is consistent with the PCA results in Fig. 4, where the first principal component (PC1, 86% of variance) is mainly associated with sequencing depth, while the biological effect appears primarily in PC2 (12% of variance).

**Table 4:** Estimated error ratio after applying normalization methods at the phylum simulation.

| Normalization | Low depth | High depth | Complete dataset |
| --- | --- | --- | --- |
| Raw data | 0.5339 | 0.2910 | 0.6720 |
| Rarefaction | 0.6261 | 0.5306 | 0.5795 |
| Centered Log Ratio | 0.4115 | 0.3811 | 0.4152 |
| Total Sum Scaling | 0.3090 | 0.4071 | 0.3565 |
| Cumulative Sum Scaling | 0.2756 | 0.3303 | 0.2763 |
| DESeq2 | 0.9660 | 0.7372 | 0.9817 |
| edgeR-TMM | 0.0556 | 0.1199 | 0.1192 |

At the phylum level, edgeR-TMM again demonstrates the lowest overall error across sequencing depths, followed by Rarefaction.

Table 4 also summarizes the effect of the normalization methods. Rarefaction and edgeR-TMM markedly reduce the error compared to raw data across depth strata, with edgeR-TMM showing the lowest overall error among all methods at the phylum level.

PCA visualizations for the phylum-level analyses are shown in Figs. 8 and 9. For edgeR-TMM normalization (Fig. 8), the score plot colored by sequencing depth (Fig. 8a) shows that sequencing depth is partially homogenized but not completely removed. The score plot colored by ecosystem (Fig. 8b) displays a clear group separation, which mainly appears in the subspace spanned by PC1 and PC3. The loading plot (Fig. 8c) reveals a relatively dispersed structure of the taxa at the phylum level. For rarefaction (Fig. 9), the score plot by sequencing depth (Fig. 9a) confirms that all normalized samples contain the same number of reads, and the score plot by ecosystem (Fig. 9b) shows that the biological effect is located in the PC1–PC2 subspace. The associated loading plot (Fig. 9c) displays a more visually structured pattern of taxa.

**Fig. 8:**
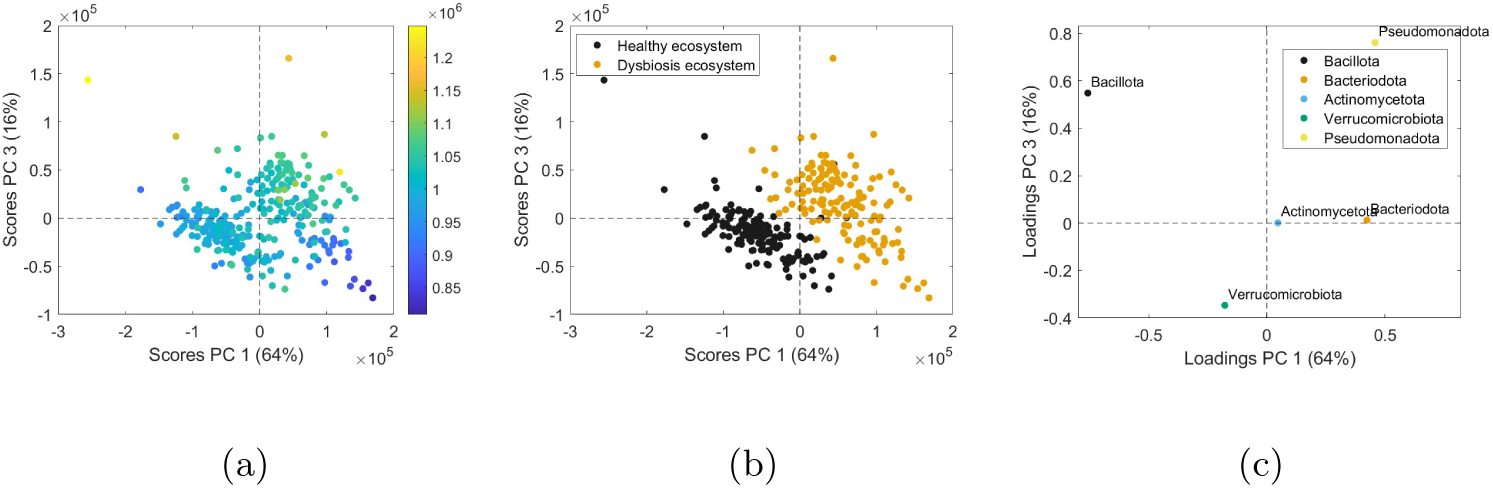
Score and loading plots of PCA for the normalized data by edgeR normalization at the phylum level: **(a)** score plot of samples colored by sequencing depth, **(b)** score plot colored by the two ecosystems, and **(c)** loading plot of bacteria at the phylum level with five taxa.

**Fig. 9:**
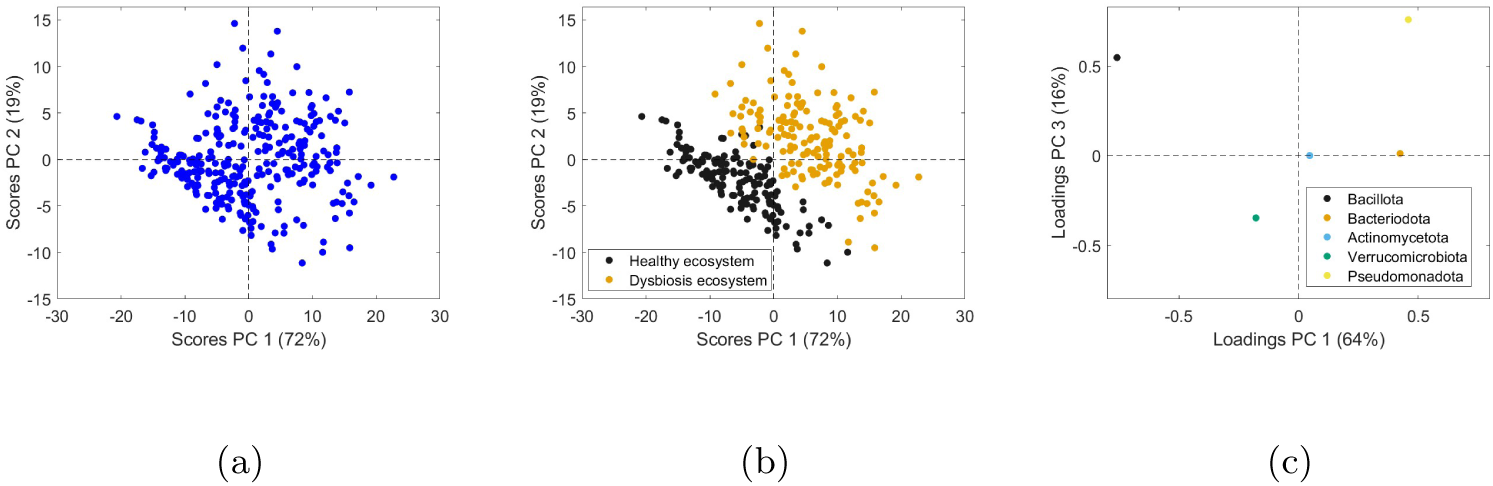
Score and loading plots of PCA for the normalized data by rarefaction at the phylum level: **(a)** score plot of samples colored by sequencing depth, **(b)** score plot colored by the two ecosystems, and **(c)** loading plot of bacteria at the phylum level with five taxa.

Table 5 presents the results from the hybrid normalization method at the phylum level, in which rarefaction is followed by a second normalization step. For some methods (e.g., DESeq2), the error is reduced after rarefaction, whereas for others (e.g., Total Sum Scaling, and edgeR-TMM) the error increases when applied on rarefied data.

**Table 5:**
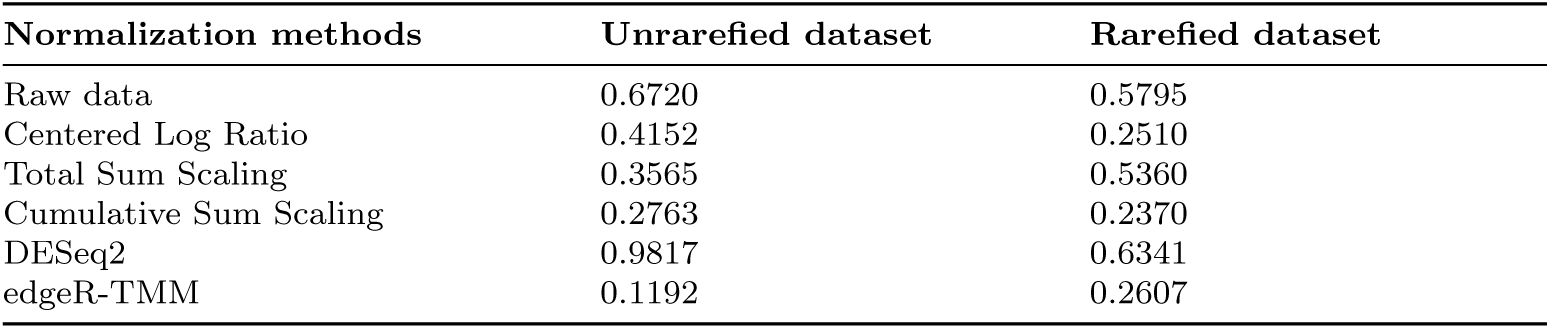
Estimated error after applying hybrid normalization model (rarefaction-based) at the phylum simulation.

At the genus level, Tables 6 and 7 report the estimated error ratios for the same set of normalization methods. In contrast to the phylum-level results, the errors at the genus level are relatively stable across low-depth, high-depth, and complete datasets. edgeR-TMM (Table 6) attains the lowest total error across all depth strata, while CLR shows the highest error. The hybrid rarefaction-based normalization (Table 7) leads to further error reductions for some methods, such as Total Sum Scaling, but not for all approaches.

**Table 6:**
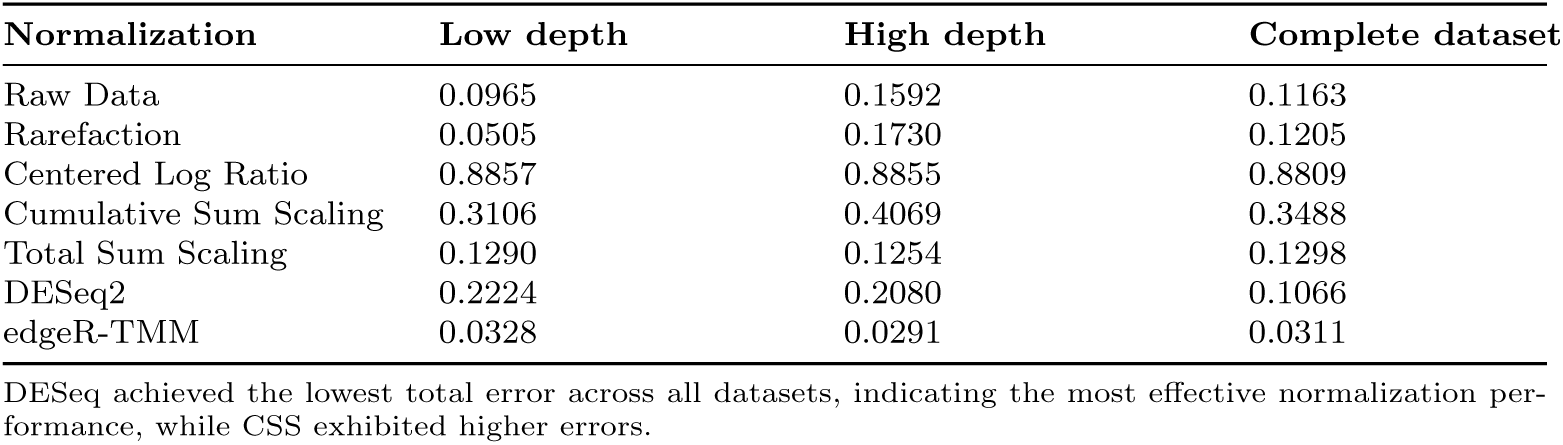
Estimated error ratio after applying normalization methods at the genus simulation.

| Normalization | Low depth | High depth | Complete dataset |
| --- | --- | --- | --- |
| Raw Data | 0.0965 | 0.1592 | 0.1163 |
| Rarefaction | 0.0505 | 0.1730 | 0.1205 |
| Centered Log Ratio | 0.8857 | 0.8855 | 0.8809 |
| Cumulative Sum Scaling | 0.3106 | 0.4069 | 0.3488 |
| Total Sum Scaling | 0.1290 | 0.1254 | 0.1298 |
| DESeq2 | 0.2224 | 0.2080 | 0.1066 |
| edgeR-TMM | 0.0328 | 0.0291 | 0.0311 |

DESeq achieved the lowest total error across all datasets, indicating the most effective normalization performance, while CSS exhibited higher errors.

**Table 7:** Estimated error after applying hybrid normalization model based on rarefaction at the genus simulation.

| Normalization methods | Unrarefied dataset | Rarefied dataset |
| --- | --- | --- |
| Raw data | 0.1613 | 0.1205 |
| Centered Log Ratio | 0.8809 | 0.9073 |
| Total Sum Scaling | 0.1298 | 0.1109 |
| Cumulative Sum Scaling | 0.3488 | 0.9073 |
| DESeq2 | 0.1066 | 0.1234 |
| edgeR-TMM | 0.0311 | 0.1215 |

Total Sum Scaling achieved the lowest total error after applying rarefaction.

PCA of genus-level data (Fig. 5.b) shows a clear separation between healthy and dysbiosis ecosystems in the raw dataset along the first principal component, where PC1 captures 82% of the variance (Fig. 5.a). After edgeR-TMM normalization (Fig. 10), the separation between ecosystems is also clearly observed along PC1, which now captures 37% of the variance. The corresponding loading and class-structure plots at the genus level are summarized in Fig. 10.

**Fig. 10:**
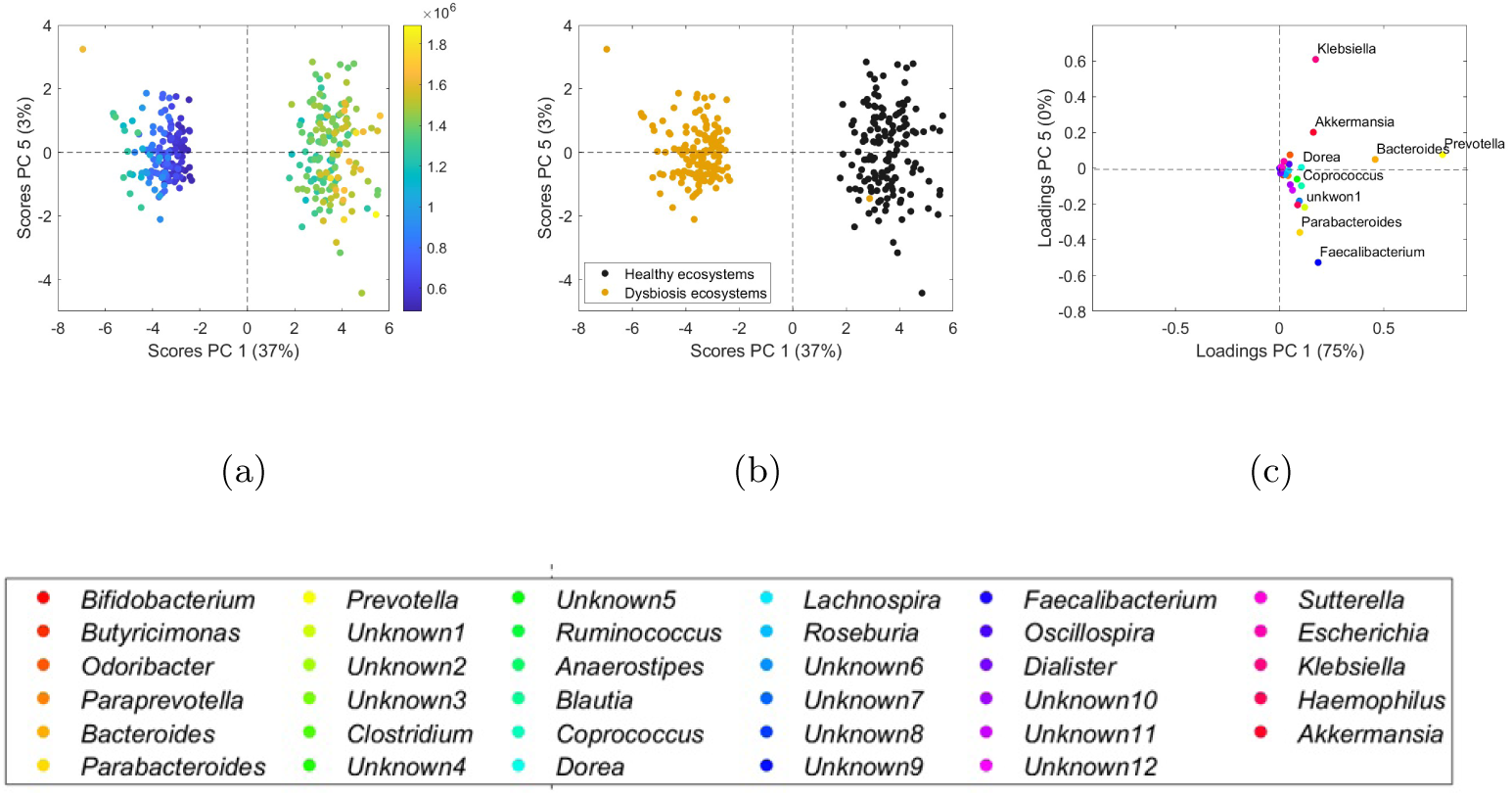
Score and loading plots of PCA for the normalized data by edgeR-TMM at the genus level: **(a)** score plot of samples colored by sequencing depth, **(b)** score plot colored by the two ecosystems, and **(c)** loading plot of bacteria at the genus level with 36 taxa. The full-width panel below shows the overall summary figure.

## 4 Discussion

At the phylum level, the results show that recovering the underlying biological signal depends both on sequencing depth and on correcting depth variability. When data are analyzed without normalization, raw datasets exhibit higher error at low depth than at high depth, while the complete dataset yields the largest error overall. This behavior arises because depth-related variability dominates the PCA structure, particularly along PC1, masking the biological effect. These observations confirm that sequencing depth acts as a confounding factor at the phylum level and motivate the need for normalization to recover biologically meaningful patterns.

In addition to the quantitative error metrics, inspection of PCA and oMEDA plots provides further insight into how normalization influences interpretation. The oMEDA errors reported in Table 4 indicate that edgeR-TMM and rarefaction are the most effective normalization methods at the phylum level, with edgeR-TMM achieving the lowest overall error. However, PCA-based representations alone can be misleading. For edgeR-TMM, the biological separation is captured in a subspace involving PC1 and PC3, resulting in a relatively complex loading structure. In contrast, rarefaction appears to remove depth-related variability more completely, with the biological effect emerging in the PC1–PC2 subspace. While this representation may seem more intuitive, oMEDA indicates that edgeR-TMM more accurately recovers the true differential abundances. This highlights that visually appealing PCA projections do not necessarily correspond to higher quantitative accuracy and underscores the value of simulation-based ground truth for objective method evaluation.

To further illustrate this point, we compared the phylum-level oMEDA plots obtained from raw data (Fig. 6.a), ground truth (Fig. 6.b), and edgeR-TMM– normalized data (Fig. 11). In the ground-truth plot, the dominant contributors are clearly identified, with *Pseudomonadota* and *Bacteriodota* associated with dysbiosis and *Bacillota* associated with healthy samples, while the remaining phyla show comparatively smaller effects. The oMEDA plot derived from raw data reproduces this structure only partially, reflecting the confounding influence of depth variability. After applying edgeR-TMM normalization, the oMEDA plot aligns more closely with the ground-truth pattern, with both the direction and relative magnitude of the main taxa more accurately recovered. In particular, *Bacteriodota*, which is largely absent in the raw-data oMEDA plot, becomes clearly detectable after edgeR-TMM normalization. Overall, this comparison shows that edgeR-TMM reduces depth-driven bias and yields oMEDA profiles that more closely reflect the expected taxa-level differences.

**Fig. 11:**
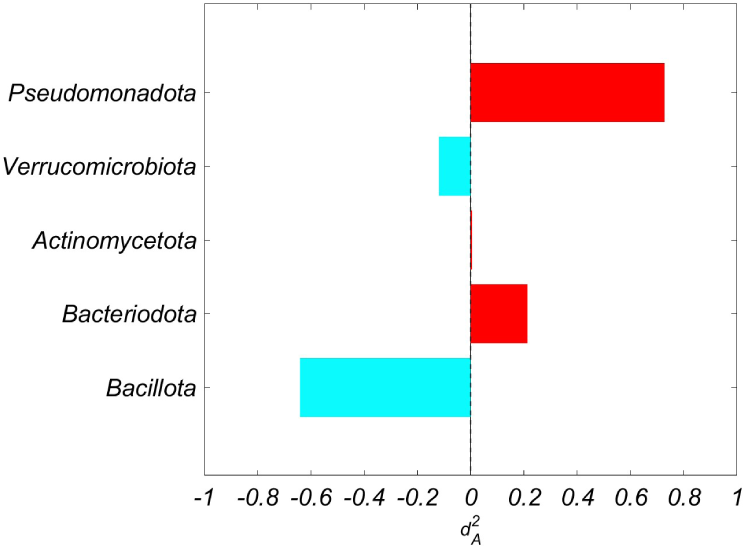
Score and comparison plots at the phylum level: Observation-based missing data methods for exploratory data analysis (oMEDA) representation of edgeR-TMM normalization.

Given the strong performance of edgeR-TMM and rarefaction at the phylum level, we further examined whether hybrid strategies could provide additional benefits. The hybrid rarefaction-based strategies (Table 5) show that combining normalization methods does not consistently improve performance. While some combinations yield modest reductions in error, others lead to poorer results, emphasizing that normalization strategies should be selected carefully rather than applied indiscriminately.

In contrast to the phylum-level results, the genus-level analysis reveals a different behavior. Errors are relatively similar across low-depth, high-depth, and complete datasets, and the PCA of raw data is already dominated by the biological effect along PC1. In this setting, sequencing depth introduces minimal depth variability, and normalization offers limited benefit. Several methods, such as CLR, substantially worsen performance, while others provide only marginal changes compared to raw data. The only exception is edgeR-TMM, which yields stable oMEDA plots and clear group separation in PCA, markedly superior to raw data according to the oMEDA error metric. This difference is further illustrated by comparing genus-level oMEDA plots obtained from raw data (Fig. 7a), ground truth (Fig. 7b), and edgeR-TMM normalization (Fig. 12). In the ground-truth plot, the biological signal is concentrated in a small number of taxa, most notably *Prevotella* and *Bacteroides*, which show strong opposing contributions between healthy and dysbiosis conditions. The raw-data oMEDA plot captures this pattern only partially, with several key taxa appearing attenuated or poorly separated from the background. After edgeR-TMM normalization, the oMEDA plot more closely resembles the ground-truth structure, with dominant taxa becoming more prominent and their relative contributions better aligned with the reference. Nevertheless, the overall improvement is less pronounced than at the phylum level, reflecting the weaker influence of depth variability in this scenario.

**Fig. 12:**
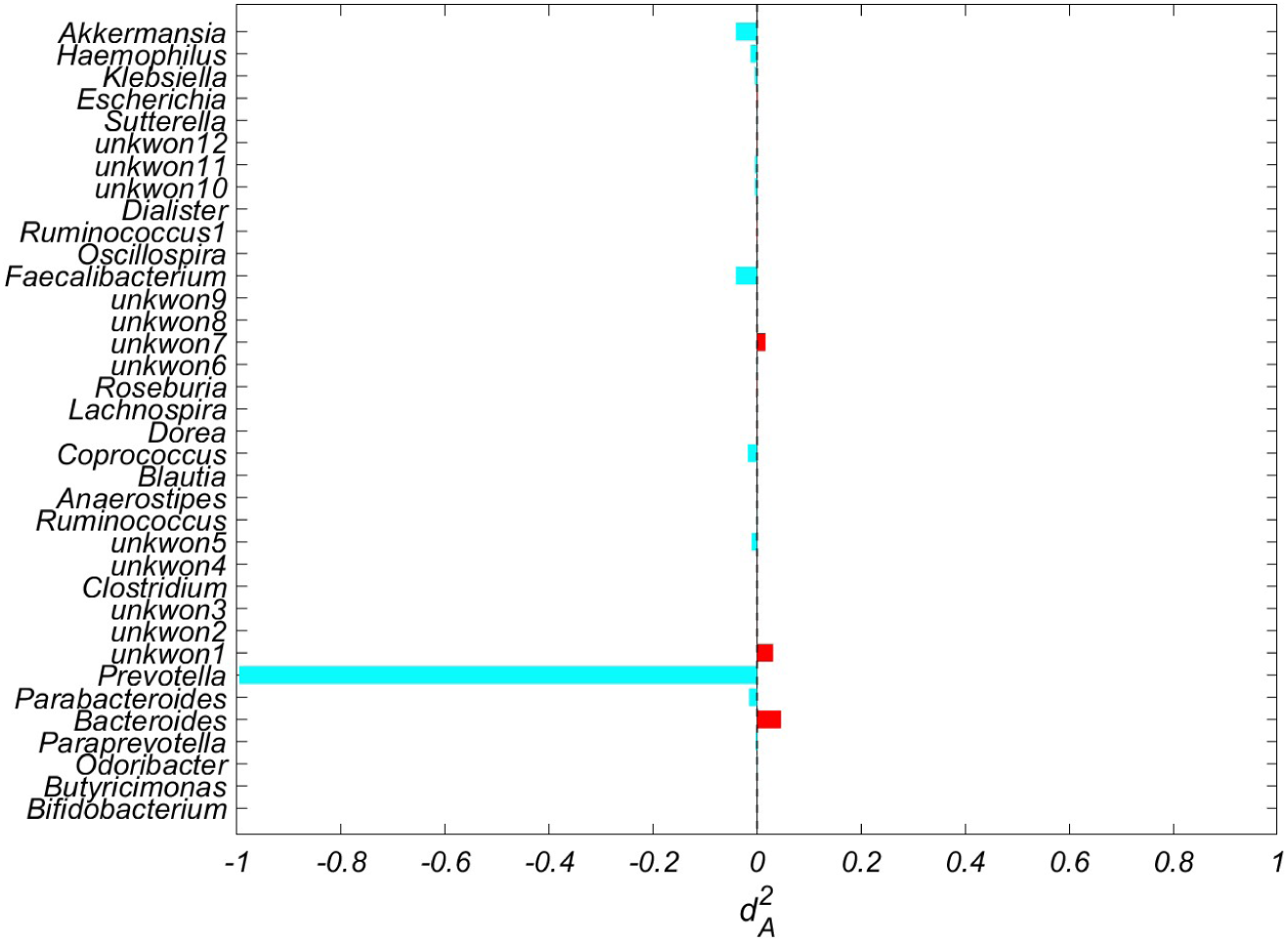
Score and comparison plots at the genus level: Observation-based missing data methods for exploratory data analysis (oMEDA) representation of edgeR-TMM normalization

These results demonstrate that the impact of normalization is highly data-dependent and varies across taxonomic resolutions. While normalization is essential at the phylum level to mitigate sequencing-depth effects, it may be unnecessary or even detrimental at the genus level when the biological signal already dominates the data structure. Combining PCA-based visualization with oMEDA and a realistic simulation framework therefore provides a more robust method for evaluating normalization methods than relying on PCA patterns alone, consistent with previous findings in microbiome studies [25, 29].

## 5 Conclusion

Our study introduces a novel simulation-based framework for the systematic evaluation of normalization methods applied to microbiome sequencing data. We assess the effectiveness of these methods simultaneously in both sample stratification and the identification of differential abundance. This assessment is performed by comparing results inferred from the normalized data using multivariate techniques against a known simulated ground-truth. The multivariate methods employed are Principal Component Analysis (PCA) and an interpretation method named observation-based Missing data methods for Exploratory Data Analysis (oMEDA). Our results demonstrate that no single approach is universally optimal, as performance varies with sequencing depth and taxonomic resolution. For example, in our case study at the phylum level, normalization using edgeR-TMM and rarefaction significantly improved accuracy by mitigating the effects of sequencing depth variation. Conversely, at the genus level, sequencing depth had less impact, and the direct analysis of raw data was sufficient, with no normalization method offering a significant improvement.

The paper also highlights a key distinction in the utility of multivariate techniques: while they are effective for inferring the accuracy of sample stratification, determining the accuracy of differential abundance necessitates knowledge of the ground-truth, making the simulation procedure instrumental. To maximize the practical utility of our framework, we recommend generating simulations that are as realistic as possible, starting from real data observations and potentially leveraging prior information from the literature in order to define (at least partially) the ground-truth. The proposed framework enables systematic evaluation of normalization methods and supports selecting the most appropriate approach for diverse microbiome datasets.

## 6 List of abbreviations

PCA (Principal Component Analysis), oMEDA (observation-based Missing data methods for Exploratory Data Analysis), OTUs (Operational Taxonomic Units), ASVs (Amplicon Sequence Variants), TSS (Total Sum Scaling), CoDA (Compositional Data Analysis), CLR (Centered Log-Ratio), PCA–oMEDA (Principal Component Analysis– observation-based Missing data methods for Exploratory Data Analysis), PCoA (Principal Coordinates Analysis), ANCOM (Analysis of Composition of Microbiomes), CSS (Cumulative Sum Scaling).

## 7 Declarations

### 7.1 Supplementary Information

The online version contains supplementary material available at https://.

- **Supplementary Material 1**
- **Supplementary Material 2**

### 7.2 Author contributions

J.C. formulated the research problem and designed the work. C.G.L. analysed the existing real dataset and generated the inputs required for the simulations. A.A. prepared the simulated data using the inputs provided by C.G.L. A.A. performed the data analysis and wrote the main manuscript text. J.C. and C.G.L. validated the analysis. A.A. prepared the figures and tables. C.G.L. and J.C. proofread and edited the manuscript. All authors have reviewed and approved the final manuscript.

### 7.3 Corresponding author

Correspondence to Amen Al Khafaji.

## 7.4 Acknowledgements

We are grateful to Alejandro Garía Vázquez for his support with programming the simulation schemes. We are also grateful to Prof. Edoardo Saccenti for his valuable guidance and insightful contributions to this study.

## 7.6 Availability of data and materials

The datasets generated during the current study and the R code used for the simulations are available in the GitHub repository: https://github.com/AmenALkhafaji/PCAoMEDA-for-assesst_norm_metageon. The code for the MEDA Toolbox is available at:https://github.com/josecamachop/MEDA-Toolbox (release v1.6). All datasets generated during the study can be reproduced using these scripts.

## 7.7 Ethics approval and consent to participate

Not applicable.

## 7.8 Consent for publication

Not applicable.

