## Supplementry file1 for "A Realistic Simulation Framework for Evaluating Microbiome Normalization in Sample Stratification and Differential Abundance"

### **Supplementary Material:** PCA Analysis and Normalization Methods for Phylum-Level Microbiome Data

#### Supplementary Discussion

This supplementary material provides detailed PCA visualizations and comparison of normalization methods at the phylum level. All figures here are referenced with an 'S' prefix (e.g., Fig. S1) to distinguish them from the main manuscript.

The PCA visualizations (Figs. 3–4) illustrate differences across normalization approaches and the raw dataset.

The Cumulative Sum Scaling (CSS) normalization (Fig. 3) produced the clearest and most distinct separation between healthy and dysbiosis groups. The corresponding

*

loading plot indicates that the *Bacillota–Bacteroidota* contrast drives the dominant axis of variation.

CLR (Fig. 1) showed good separation, slightly less distinct than CSS, but retained the primary biological gradient. TSS (Fig. 2) produced moderate separation with potential overlap, while DESeq (Fig. 4) yielded poor separation and high overlap. The raw unnormalized data confirmed the need for normalization.

Comparison with the simplified directional vector confirms that the primary gradient corresponds to the *Bacillota–Bacteroidota* trade-off. Minor discrepancies for *Pseudomonadota*, *Verrucomicrobiota*, and *Actinomycetota* highlight secondary contributions.

Overall, CSS and CLR are most suitable for phylum-level analyses in terms of visual separation and alignment with oMEDA results (Table 4 in the main text). TSS and DESeq show weaker correction of technical variability.

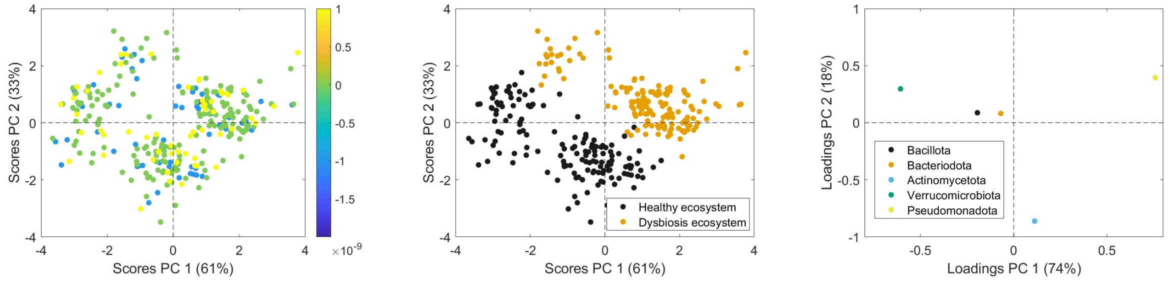


(a) (b) (c)

Figure 1: Score and loading plots of PCA for the normalized data by Centred Log Ratio (CLR) normalization at the phylum level: **(a)** score plot of samples colored by sequencing depth, **(b)** score plot colored by the two ecosystems, and **(c)** loading plot of bacteria at the phylum level with five taxa.

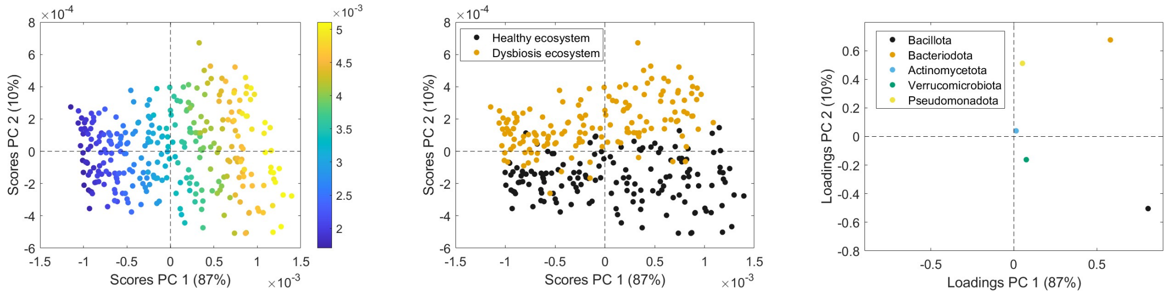


(a) (b) (c)

Figure 2: Score and loading plots of PCA for the normalized data by Total Sum Scale (TSS) normalization at the phylum level: **(a)** score plot of samples colored by sequencing depth, **(b)** score plot colored by the two ecosystems, and **(c)** loading plot of bacteria at the phylum level with five taxa.

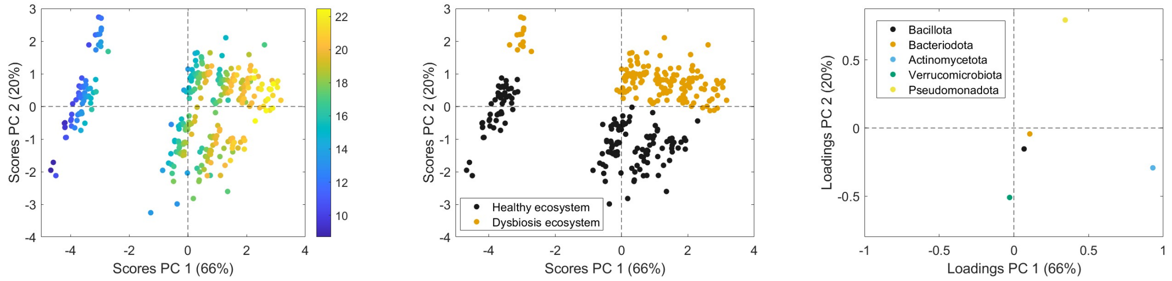


**(a)** **(b)** **(c)**

Figure 3: Score and loading plots of PCA for the normalized data by Cumulative Sum Scaling (CSS) normalization at the phylum level: **(a)** score plot of samples colored by sequencing depth, **(b)** score plot colored by the two ecosystems, and **(c)** loading plot of bacteria at the phylum level with five taxa.

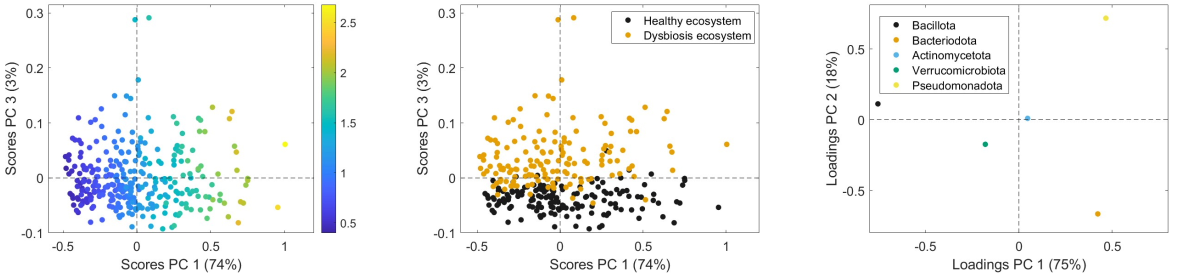


**(a)** **(b)** **(c)**

Figure 4: Score and loading plots of PCA for the normalized data by DESeq normalization at the phylum level: **(a)** score plot of samples colored by sequencing depth, **(b)** score plot colored by the two ecosystems, and **(c)** loading plot of bacteria at the phylum level with five taxa.
