## Supplementry file2 for "A Realistic Simulation Framework for Evaluating Microbiome Normalization in Sample Stratification and Differential Abundance"

### **Supplementary Material:** PCA Analysis and Normalization Methods for Genus-Level Microbiome Data

#### Supplementary Discussion

This supplementary material provides an overview of PCA visualizations and normalization methods at the genus level. Figures are referenced using Fig. 1-6.

Raw data often show the strongest natural separation between groups because the original abundance levels are preserved. Rarefaction can reduce this separation by discarding reads to equalize depth. Transformations like CLR, TSS, and CSS adjust for compositional effects, but may compress differences between groups depending on the data structure. In contrast, differential-abundance methods such as EdgeR model count

*

distributions directly, and can detect group differences statistically even when visual separation in transformed data appears weaker (see Figs. 4 and 6).

Across the depth-related datasets, Rarefaction shows the clearest separation between the healthy and dysbiosis groups. CLR and TSS produce tight score ranges that can cause overlap, while CSS captures little variation and separates groups poorly. EdgeR shows high variance but is affected by extreme scaling, making the separation less reliable. The raw gen data also separates groups well, but remains influenced by depth differences. In contrast, Rarefaction provides both high variance and stable scaling, making it the most effective method for visualizing group distance under depth variation (see Figs. 1–4).

When comparing PC1 loadings to the biological effect sizes, the gen method shows the strongest overall agreement. Its high PC1 variance and wide loading range allow the major genera with large effect sizes, such as Prevotella, Bacteroides, and Oscillospira, to appear more prominently along PC1, making their influence easier to detect. CLR also performs reasonably well, maintaining good variance and preserving relative differences between taxa, which helps it capture part of the biological pattern even if the loadings are more compressed. In contrast, CSS explains less variation, and therefore aligns weakly with the true effect sizes. The methods that provide only genus lists, such as EdgeR, Rarefaction, and TSS, cannot be evaluated quantitatively, so their agreement with effect sizes remains unclear. Overall, gen offers the closest match to the biological signals in your effect size data, with CLR being the most dependable among the normalized options (Figs. 1–6).

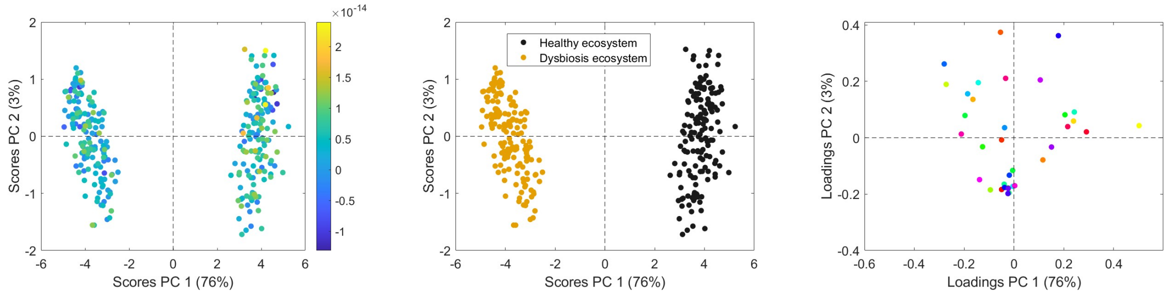


(a) (b) (c)

Figure 1: PCA visualization for Centred Log Ratio (CLR) normalization at the genus level. Each panel shows different aspects of the data: (a) colored by sequencing depth, (b) colored by the two ecosystems, (c) genus-level loadings with 36 taxa.

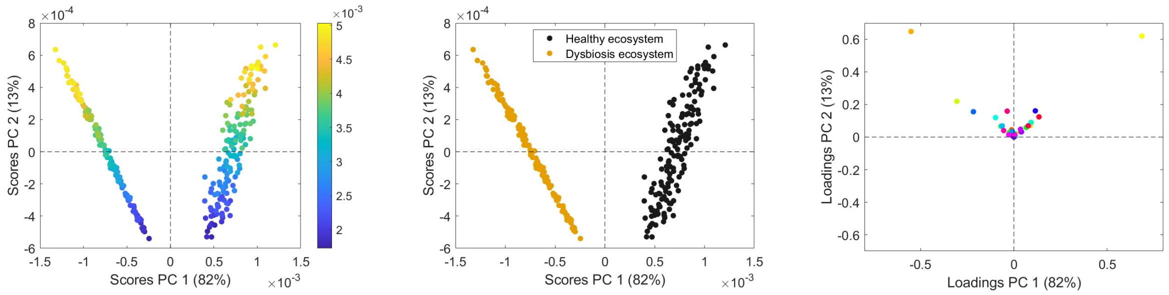


(a) (b) (c)

Figure 2: PCA visualization for Total Sum Scale (TSS) normalization at the genus level. Each panel shows different aspects of the data: (a) colored by sequencing depth, (b) colored by the two ecosystems, (c) genus-level loadings with 36 taxa.

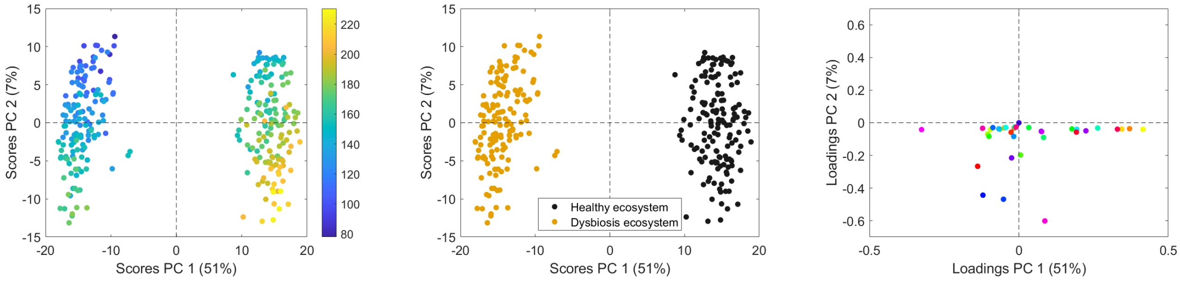


(a) (b) (c)

Figure 3: PCA visualization for Cumulative Sum Scaling (CSS) normalization at the genus level. Each panel shows different aspects of the data: (a) colored by sequencing depth, (b) colored by the two ecosystems, (c) genus-level loadings with 36 taxa.

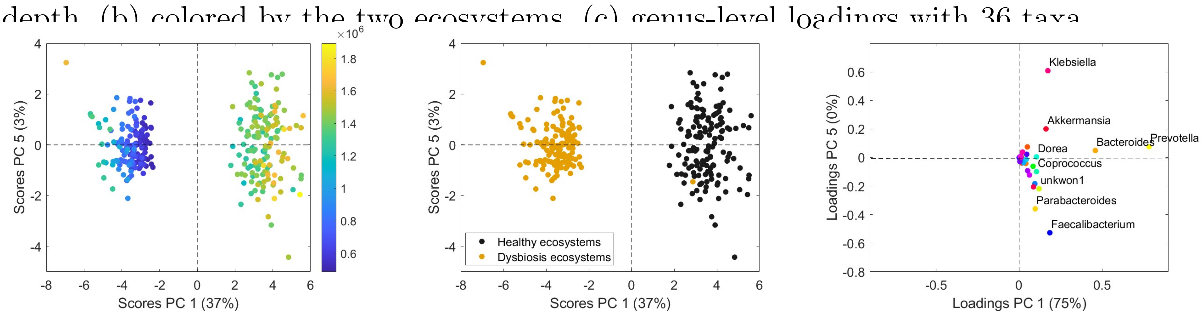


(a) (b) (c)

Figure 4: PCA visualization for EdgeR normalization at the genus level. Each panel shows different aspects of the data: (a) colored by sequencing depth, (b) colored by the two ecosystems, (c) genus-level loadings with 36 taxa.

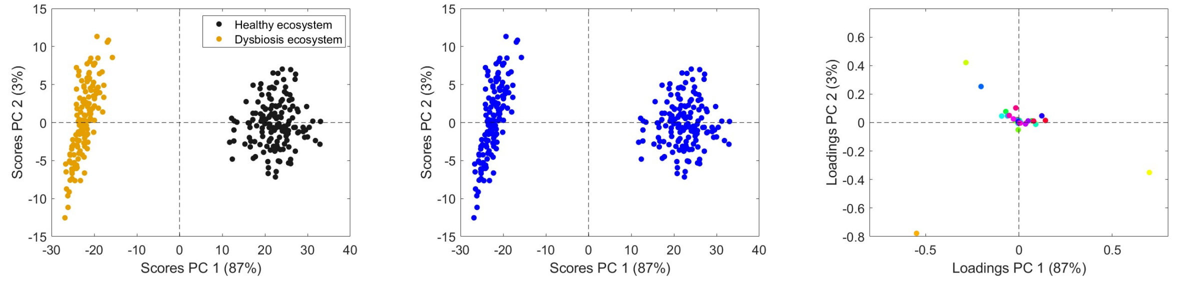


(a) (b) (c)

Figure 5: PCA visualization for Rarefaction normalization at the genus level. Each panel shows different aspects of the data: (a) colored by sequencing depth, (b) colored by the two ecosystems, (c) genus-level loadings with 36 taxa.

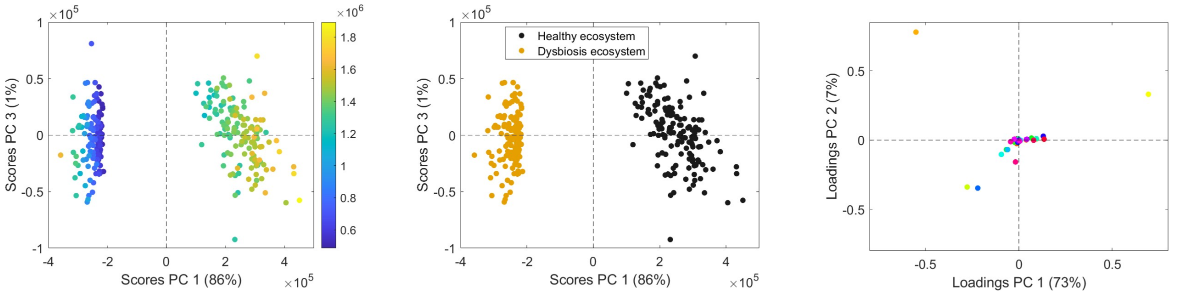


(a) (b) (c)

Figure 6: PCA visualization for DESeq normalization at the genus level. Each panel shows different aspects of the data: (a) colored by sequencing depth, (b) colored by the two ecosystems, (c) genus-level loadings with 36 taxa.
