## Supplementry file3 for "A Realistic Simulation Framework for Evaluating Microbiome Normalization in Sample Stratification and Differential Abundance"

### **Supplementary Material:** PCA Analysis and Hybrid Normalization Methods for Phylum-Level Microbiome Data

#### Supplementary Discussion

The comparison of hybrid normalization methods for the phylum-level data reveals clear differences in their performance across three main aspects: separation of healthy and dysbiosis ecosystems, control of sequencing depth effects, and agreement of taxa loadings with known biology. The PCA visualizations provide an intuitive representation of these effects, while the oMEDA error metrics allow quantitative assessment of their accuracy.

CSS and DESeq normalization produce the most pronounced separation between healthy and dysbiosis samples along the primary principal component, as illustrated in Figs. 3 and 4. TSS shows moderate separation (Fig. 2), whereas CLR and edgeR-TMM

*

produce clear but slightly less dramatic separation (Figs. 1 and 5). Raw data exhibits weak separation, with substantial overlap between groups. These patterns indicate that CSS and DESeq are highly effective at isolating the primary biological signal from technical variability.

Evaluation of depth variation using the score plots highlights differences in the ability of each method to mitigate sequencing depth effects. CSS and DESeq achieve the tightest clustering of samples, suggesting effective removal of technical variation. CLR, TSS, and edgeR-TMM display moderate spread along both axes, indicating residual depth-related variability. Raw data shows extreme dispersion of sample scores, reflecting poor handling of library size differences.

Comparison of PCA loadings with the simplified biological vector (Bacillota: 5, Bacteroidota: -5, Actinomycetota: -0.45, Verrucomicrobiota: 3, Pseudomonadota: -8) reveals that CLR and edgeR-TMM faithfully capture the primary biological gradient, while CSS, DESeq, and TSS may contradict expected biological patterns.

Overall, normalization effects are context-dependent. CLR normalization emerges as a particularly suitable approach for phylum-level data when the goal is to recover biologically meaningful patterns while maintaining reasonable control of sequencing depth. The full set of PCA score and loading plots supporting this comparison is presented in Figs. 1–5.

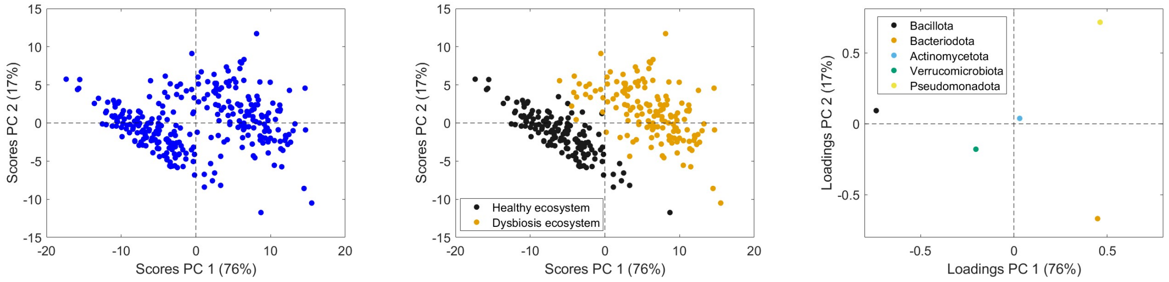


(a) (b) (c)

Figure 1: Score and loading plots of PCA for Centered Log Ratio (CLR) normalization at the phylum level: (a) score plot of samples colored by sequencing depth, (b) score plot colored by the two ecosystems, (c) loading plot of bacteria at the phylum level with five taxa.

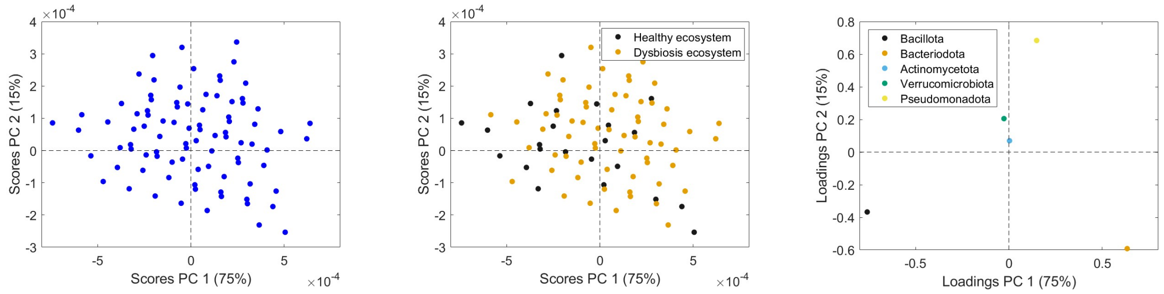


(a) (b) (c)

Figure 2: Score and loading plots of PCA for Total Sum Scale (TSS) normalization at the phylum level: (a) score plot of samples colored by sequencing depth, (b) score plot colored by the two ecosystems, (c) loading plot of bacteria at the phylum level with five taxa.

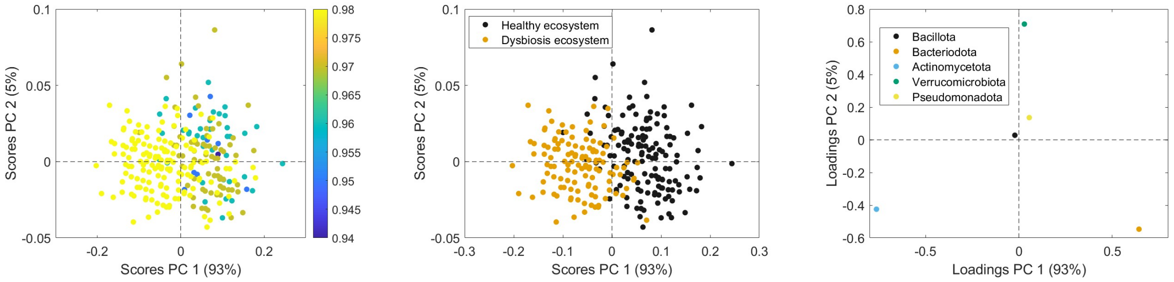


(a) (b) (c)

Figure 3: Score and loading plots of PCA for Cumulative Sum Scaling (CSS) normalization at the phylum level: (a) score plot of samples colored by sequencing depth, (b) score plot colored by the two ecosystems, (c) loading plot of bacteria at the phylum level with five taxa.

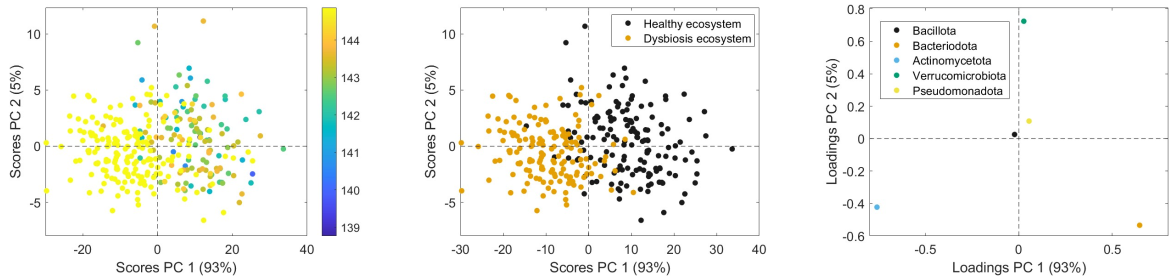


(a) (b) (c)

Figure 4: Score and loading plots of PCA for DESeq2 normalization at the phylum level: (a) score plot of samples colored by sequencing depth, (b) score plot colored by the two ecosystems, (c) loading plot of bacteria at the phylum level with five taxa.

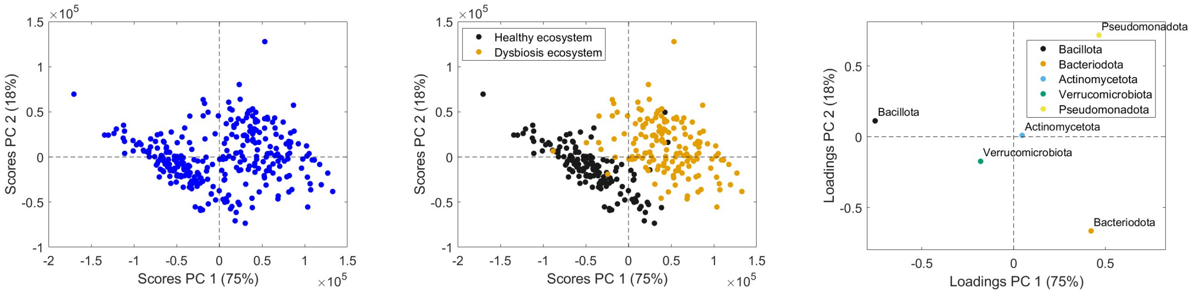


(a) (b) (c)

Figure 5: Score and loading plots of PCA for edgeR-TMM normalization at the phylum level: (a) score plot of samples colored by sequencing depth, (b) score plot colored by the two ecosystems, (c) loading plot of bacteria at the phylum level with five taxa.
