## Supplementry file4 for "A Realistic Simulation Framework for Evaluating Microbiome Normalization in Sample Stratification and Differential Abundance"

### **Supplementary Material:** PCA Analysis and Hybrid Normalization Methods for Genus-Level Microbiome Data

#### Supplementary Discussion

This supplementary material provides an overview of PCA visualizations and hybrid normalization methods at the genus level. Figures are referenced using Fig. 1-5.

Based on the PCA plots from various hybrid normalization methods, raw data (non-normalized) shows the clearest visual distinction between the healthy ecosystem (black group) and the dysbiosis ecosystem (yellow group, see Fig. 6). CLR and CSS exhibit low PC1 variance and very narrow score ranges, resulting in tightly clustered points with substantial overlap. DESeq2 improves separation somewhat, with moderate PC1 variance and a reasonable score range, though some overlap remains. EdgeR shows high variance,

*

but scores are skewed and on a very large scale, potentially distorting group boundaries. TSS captures high variance, but extremely small scores lead to tight clustering and overlap. In contrast, raw data provides a wide spread along PC1, allowing distinct clustering between healthy and dysbiosis ecosystems. While non-normalized data can be influenced by technical factors such as sequencing depth, it provides the best visual distinction. For analyses requiring normalization, DESeq2 offers a reasonable alternative, maintaining moderate separation and variance. Based on the current plots, raw data remains optimal for assessing visual group distance, although hybrid approaches combining normalization and rarefaction could be explored.

When considering depth variation, raw data exhibits wide variation along PC1, allowing clear separation between groups, but this separation is strongly influenced by depth-related variation. CLR and CSS show very low PC1 variance and narrow score ranges, effectively controlling depth variation but at the cost of biological signal and group separation. DESeq2 provides a better compromise, maintaining separation while substantially reducing depth influence. EdgeR captures high variance, but the large score range suggests that some separation may be driven by technical artifacts. TSS shows high variance, but very small scores again limit separation. Overall, DESeq2 offers the optimal balance between group separation and control of depth-related variation.

Based on loadings data from various normalization methods and effect sizes for genera, raw data shows the strongest alignment with biological variation. Its PC1 captures most of the variance and exhibits a wide loading range, allowing major genera with large effect sizes, such as Prevotella, Bacteroides, and Oscillospira, to contribute prominently. DESeq2 also performs reasonably well, though slightly inferior due to lower variance and narrower loadings. EdgeR has high PC1 variance, but the narrow loading range may not adequately reflect large effect sizes. CLR and CSS show low variance and small loadings, limiting their ability to highlight strong-effect genera. TSS provides only genus lists without numerical loadings, preventing quantitative evaluation. Overall, raw data is the preferred method for analyses focused on genus contributions, while DESeq2 is a reasonable alternative when normalization is required (see Figs. 1-5).

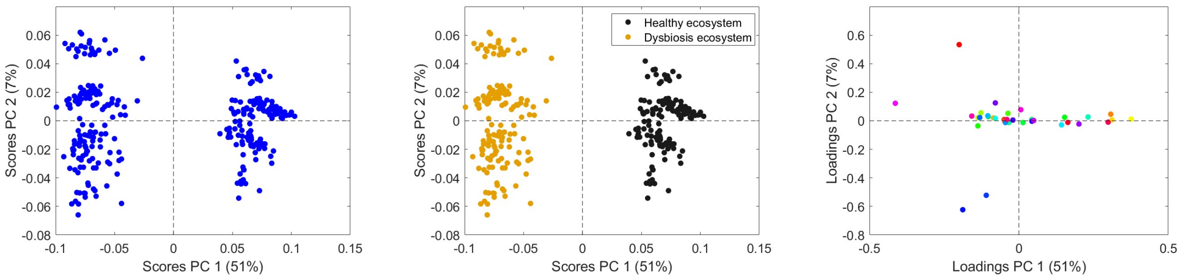


(a) (b) (c)

Figure 1: Score and loading plots of PCA for Centered Log Ratio (CLR) normalization at the genus level: (a) score plot of samples colored by sequencing depth, (b) score plot colored by the two ecosystems, (c) loading plot of bacteria at the genus level.

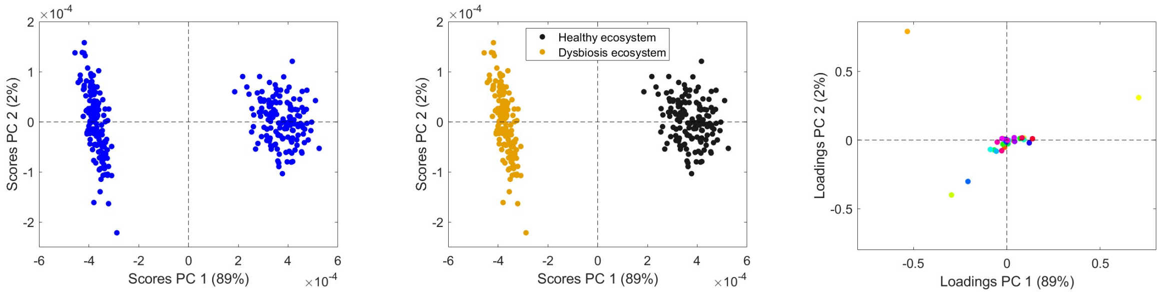


(a) (b) (c)

Figure 2: Score and loading plots of PCA for Total Sum Scale (TSS) normalization at the genus level: (a) score plot of samples colored by sequencing depth, (b) score plot colored by the two ecosystems, (c) loading plot of bacteria at the genus level.

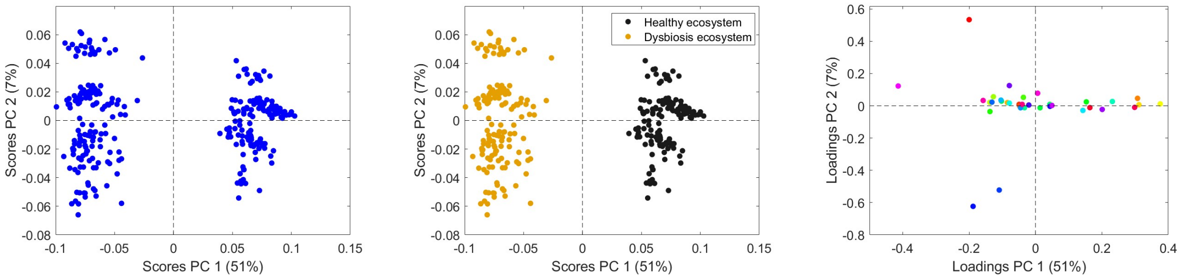


(a) (b) (c)

Figure 3: Score and loading plots of PCA for Cumulative Sum Scaling (CSS) normalization at the genus level: (a) score plot of samples colored by sequencing depth, (b) score plot colored by the two ecosystems, (c) loading plot of bacteria at the genus level.

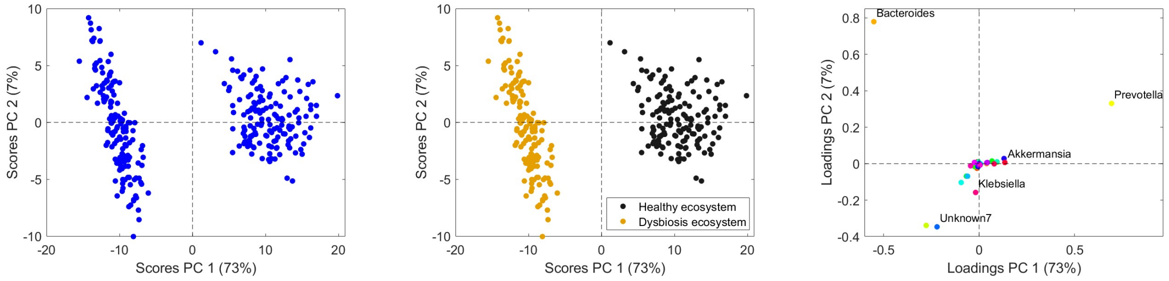


(a) (b) (c)

Figure 4: Score and loading plots of PCA for DESeq2 normalization at the genus level: (a) score plot of samples colored by sequencing depth, (b) score plot colored by the two ecosystems, (c) loading plot of bacteria at the genus level.

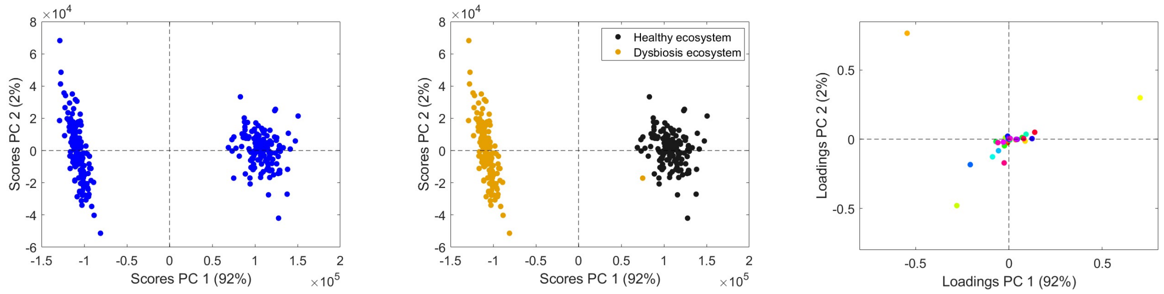


(a) (b) (c)

Figure 5: Score and loading plots of PCA for EdgeR normalization at the genus level: (a) score plot of samples colored by sequencing depth, (b) score plot colored by the two ecosystems, (c) loading plot of bacteria at the genus level.

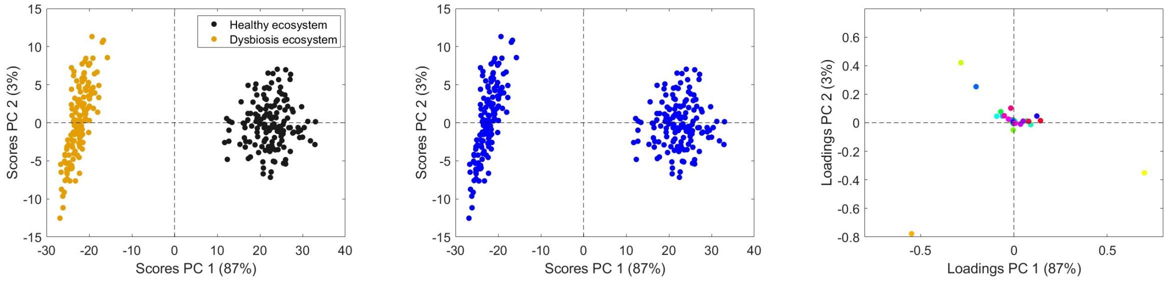


(a) (b) (c)

Figure 6: Score and loading plots of PCA for Rarefaction normalization at the genus level: (a) score plot of samples colored by sequencing depth, (b) score plot colored by the two ecosystems, (c) loading plot of bacteria at the genus level.
